# Early to Middle Pleistocene transition shapes the evolution of human-specific mutations associated with height and basal metabolic rate

**DOI:** 10.1101/2024.06.23.600148

**Authors:** Yufeng Zhang, Jie Wang, Chuanyou Yi, Yue Su, Zi Yin, Shuxian Zhang, Ke Wang, He Huang, Jin Li, Shaohua Fan

## Abstract

Understanding the genetic basis of modern-human-specific traits is essential for elucidating the formation of anatomically modern humans (AMHs). Here, we studied the genetic underpinnings of height and basal metabolic rate (BMR), which have undergone extensive modifications in AMHs compared to other *Homo* species and apes. The results revealed a significant genetic correlation between the two traits. The evolution of the variants associated with height and BMR was heavily influenced by environmental factors, marked by two bursts during the Early to Middle Pleistocene transition and one afterward, accounting for 37.4% of the inferred causal variants for height and BMR. We identified an AMH-specific mutation, rs34590044-A, which emerged around 981,916 years ago, coinciding with the first burst of variants associated with increased height and BMR. rs34590044-A upregulates the expression of *ACSF3* via increasing its enhancer activity, leading to increased mitochondrial function, body length, and BMR exclusively in mice fed essential amino acids, specifically threonine-enriched diets, which are characteristic of meat-based diets. Therefore, the emergence of rs34590044-A may contribute to the shift from an herbivorous to a carnivorous diet in AMHs. Our results underscore the complex interplay between genetics and environment in shaping the crucial phenotypes and physiological traits of AMHs.

## Introduction

Over the past 6–15 million years, as the human lineage diverged from our closest living relatives, chimpanzees and other great apes, the ancestors of modern humans underwent remarkable transformations, culminating in the emergence of anatomically modern humans (AMHs)^1–3^. For example, previous studies have revealed that modern humans possess a basal metabolic rate (BMR) three times higher than that of non-human apes, alongside a general trend towards increased stature, from *Australopithecus* to *Homo erectus*, which mirrors the heights of AMHs^4^.

It has been posited that a dietary shift from herbivory to carnivory may have played a pivotal role in the evolution of traits in AMHs, including increased height and BMR, alongside other notable characteristics such as the development of larger, more complex brains, bipedalism, an extended lifespan, reduced gut length, smaller yet sharper teeth, and augmented social cohesion^5–7^. This hypothesis stems from the recognition that meat-enriched diets offer a concentrated source of nutrients, including all essential amino acids (EAAs), bioavailable iron and zinc, and essential vitamins^8^. Additionally, meat consumption provides omega-3 fatty acids that are vital for brain development, and fat stores that boast more than double the energy density per gram compared to plant-based food^8^. However, the mechanism and genetic basis of this transition remain elusive. Furthermore, it is unclear whether the increased BMR and height in AMH were caused by the pleiotropic effect of the same mutation or two independent adaptation processes.

In this study, we uncovered a common genetic foundation underlying both human height and BMR through genome-wide association analysis of 458,303 UK Biobank participants^9^. We identified 6,385 shared candidate causal variants, accounting for 19% of the candidate causal variants of height and 29% of candidate causal variants of BMR. Our results indicate that environmental factors played a pivotal role in shaping the emergence of height and BMR-associated variants, with two distinct bursts of such variants occurring during the Early to Middle Pleistocene transition (MPT) and one additional burst occurring after the transition.

In addition, we identified an AMH-specific mutation, rs34590044-A, which is associated with increased stature and elevated BMR in humans. rs34590044-A emerged approximately 981,916 years ago, coinciding with the first burst of the variants associated with height and BMR. rs34590044-A is located in an enhancer region of Acyl-CoA Synthetase Family Member 3 (*ACSF3*). Some rare mutations in the coding region of *ACSF3* have been reported to be associated with the suppressed mitochondrial activity and accumulation of the toxic metabolites methylmalonate (MMA) and malonate, leading to a disease called combined malonic and methylmalonic aciduria (CMAMMA) in humans^10–12^. However, these rare mutations cannot explain the observations in previous studies that *ACSF3* is expressed at a higher level in humans compared to chimpanzees, gorillas, and orangutans^13,14^.

Our results suggest that rs34590044-A is significantly associated with the elevated enhancer activity and gene expression of *ACSF3*. In addition, *in vitro*, *ex vivo*, and *in vivo* experiments demonstrated that the elevation of ACSF3 enhances mitochondrial activity and mitigates the accumulation of the toxic MMA byproduct, which likely contributes to significant increases in body length and energy expenditure in mice. Particularly, the increased body length and energy expenditure were observed exclusively in mice fed EAAs in specifically threonine-enriched diets that were characteristic of meat-based diets. Therefore, the emergence of rs34590044-A is likely contributed to the transition from an herbivorous to a carnivorous diet in AMHs as well as their increased height and BMR. Our findings offer crucial insights into how the intricate interaction between genetic and environmental factors shapes the evolution of key phenotypic and physiological traits within the human species.

## Results

### Genetic correlation between height and BMR

To explore the potential interplay between height and BMR, we first calculated the genetic correlation through cross-trait linkage disequilibrium (LD) score regression method^15^ between these two traits based on the genome-wide association study (GWAS) summary statistics of 458,303 UK Biobank (UKBB) samples with a linear mixed model^9^. We observed a significant positive correlation between height and BMR (r_g_ = 0.6108, standard error = 0.01482, *P* < 1 × 10^-5^), which aligned well with previous reports^16^ (r_g_ = 0.61, standard error = 0.0148, *P* = 1 × 10^-308^). This result suggests that shared genetic factors influence both height and BMR.

To explore genetic underpinnings of height and BMR at a finer scale, we calculated the local genetic correlations between the two traits using Local Analysis of Variant Association (LAVA)^17^ across 2,495 non-overlapping genomic loci (∼1 Mb each) with minimal LD between them. First, we performed univariate tests to filter out loci lacking significant genetic signals for either trait, as these loci cannot yield reliable local genetic correlations^17^. In total, 1,499 loci (60.08%) exceeded genome-wide significance for associations with both height and BMR (*P* < 2 × 10^-5^; Bonferroni correction threshold of 0.05/2,495). Next, we calculated local genetic correlations between these traits for each of the 1,499 loci using bivariate LAVA. A substantial proportion of loci (33.89%, or 508 out of 1,499 loci) demonstrated significant local genetic correlations after Bonferroni correction (*P* < 3.34 × 10^-5^; 0.05/1,499) (**Fig. 1a and Supplementary Table 1**). Notably, all 508 significant local correlations were positive, ranging from 0.29 to 0.94, with average and median correlation values of 0.62 and 0.60, respectively (**Fig. 1b**). These results align closely with the observation of a global genetic correlation of 0.61 between height and BMR estimated across the whole genome.

**Fig 1.**
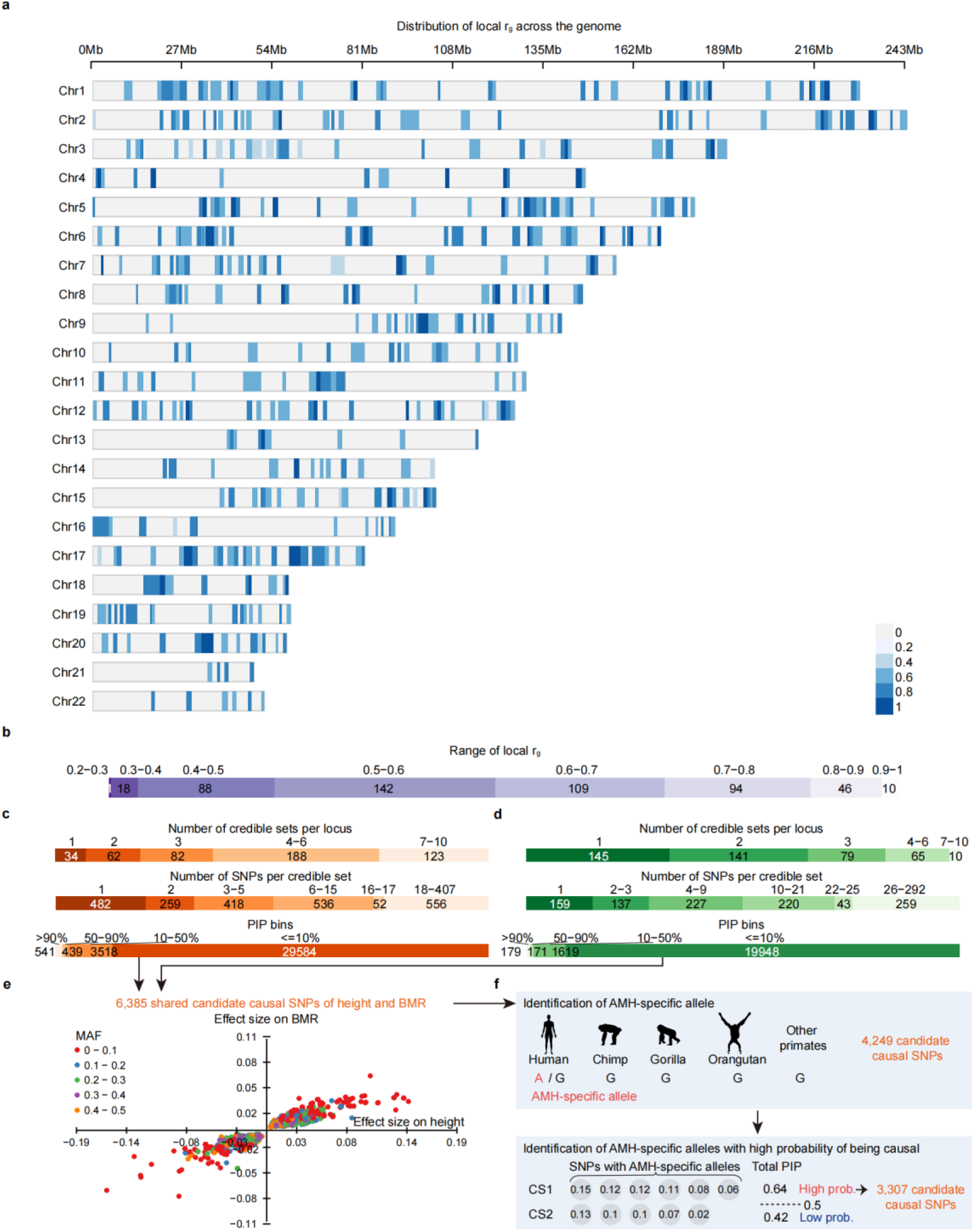
Significant genetic correlation between height and basal metabolic rate (BMR) **a**: Distribution of the significant local rg of height and BMR at 508 loci in the human genome (GRCh37). Each bar represents a locus, and the color indicates the level of genetic correlation. **b**: Number of loci binned based on the local rg. **c-d**: Summaries of fine-mapped credible sets (CSs) and SNPs for height (**c**) and BMR (**d**). For each trait, the number of independent 95% CSs per region, the number of fine-mapped SNPs per 95% CS and the number of fine-mapped SNPs binned based on posterior inclusion probability (PIP) are shown. **e**: Comparison of the effect size of 6,385 candidate causal SNPs for height (x-axis) and BMR (y-axis). Each dot is a SNP and the color indicates minor allele frequency (MAF). **f**: Schematic demonstration showing the identification of candidate causal SNPs with anatomically modern humans (AMH)-specific alleles and those exhibiting high probability of being causal (greater than 50% per CS).. The silhouettes are described in **Methods**.

### Shared genetic basis of height and BMR

We next performed statistical fine-mapping using SuSiE^18^ to detect the underlying shared causal genetic single nucleotide polymorphisms (SNPs) between height and BMR. We excluded fourteen loci, including MHC regions and two additional long-range LD regions (chr11 48-50M; GRCh37) (**Supplementary Table 2**), which often lead to false positives for SuSiE and other fine-mapping software^19^. We then performed statistical fine-mapping at the remaining 494 loci for height and BMR using SuSiE (**Methods**).

Our study detected 34,082 potentially causal SNPs associated with height across 489 out of 494 loci (**Fig. 1c and Supplementary Table 3**). These SNPs are dispersed among 2,303 95% credible sets (95% CS), indicating the presence of at least 2,303 causal SNPs contributing to height variation within the sample population^20^. Similarly, we identified 21,917 potentially causal SNPs of BMR among 1,045 95% CSs located at 440 out of 494 loci, suggesting that 1,045 causal SNPs influence BMR (**Fig. 1d and Supplementary Table 4**). Notably, 93.05% (455) of loci exhibited multiple CSs for height, in contrast to 67.05% (295) of loci for BMR (**Figs. 1c and 1d**), implying a potentially more complex polygenic architecture underlying height variation.

We found 6,385 shared potentially causal SNPs for height and BMR across 342 loci (within 550 CSs and 547 CSs based on the fine-mapping of height and BMR, respectively) (**Supplementary Table 5**), accounting for 19% of the causal SNPs of height and 29% of the causal SNPs of BMR. Notably, all shared causal SNPs for both height and BMR displayed consistent effect directions of both traits, aligning with the strong positive local genetic correlations (**Fig. 1e**).

Interestingly, 66.55% (4,249 SNPs within 500 CSs located in 328 loci, based on the fine-mapping of height) of these shared causal SNPs harbored at least one AMH-specific allele, which are absent in the genomes of 26 non-AMH primates (**Fig.1f and Methods**). Moreover, 77.83% (3,307 SNPs located in 261 loci, based on the fine-mapping of height) of SNPs with AMH-specific alleles exhibited a cumulative posterior inclusion probability (PIP) exceeding 50% in 72% (360 out of 500, based on the fine-mapping of height) of CSs (**Fig.1f, Supplementary Table 5 and Methods**), suggesting a high likelihood that these SNPs harboring AMH alleles are causal to height and BMR. This observation underscores the potential impact of AMH-specific alleles on the evolution of distinctive human traits and prompts further investigation into their evolutionary history and functional significance.

### Co-evolution of height and BMR

Based on Relate^21^, a method used to infer the age of alleles based on genome-wide genealogies, we obtained the allele ages of 3,166 (95.74%) of the 3,307 shared potential causal SNPs for both height and BMR with AMH-specific alleles across 26 global populations in the 1000 Genomes project^22^ (1KGP), among which 3,154 SNPs whose ages for derived alleles were estimated (**Supplementary Table 6**).

We observed that the ages of the potential causal SNPs for height and BMR ranged from 4.31 million years ago to 5,092 years ago, with a median age of 759,478 years ago, indicating the dynamic evolution of height and BMR in the present-day British (GBR) population. These 3,150 age-estimated SNPs were separated into two groups based on the effects of AMH-specific alleles on height and BMR (**Fig. 2a**). Of these, 52.2% (1,645) exhibited AMH-specific alleles associated with increased height and BMR, while the remaining 47.8% (1,505) of the SNPs carried AMH-specific alleles associated with decreased height and BMR in the UKBB. In addition, the ages of AMH-specific alleles with decreased effects were significantly older than those with increased effects, with a median age of 811,020 years for AMH-specific alleles with decreased effects, compared to 696,168 years for AMH-specific alleles with increased effects (one-tailed Wilcoxon rank sum test, *P* = 1 × 10^-4^) (**Fig. 2b**). The same observation was also significant in the other 18 populations in the 1KGP (one-tailed Wilcoxon rank sum test, Benjamini–Hochberg corrected *P* < 0.05) (**Extended Data Fig. 1 and Supplementary Table 7**). Although the remaining seven populations did not reach statistical significance (Benjamini– Hochberg corrected *P* > 0.05, **Supplementary Table 7**), the median ages of the AMH-specific alleles associated with decreased height and BMR were older than the ones associated with increased height and BMR in all the seven populations (**Extended Data Fig. 1 and Supplementary Table 7**).

**Fig 2.**
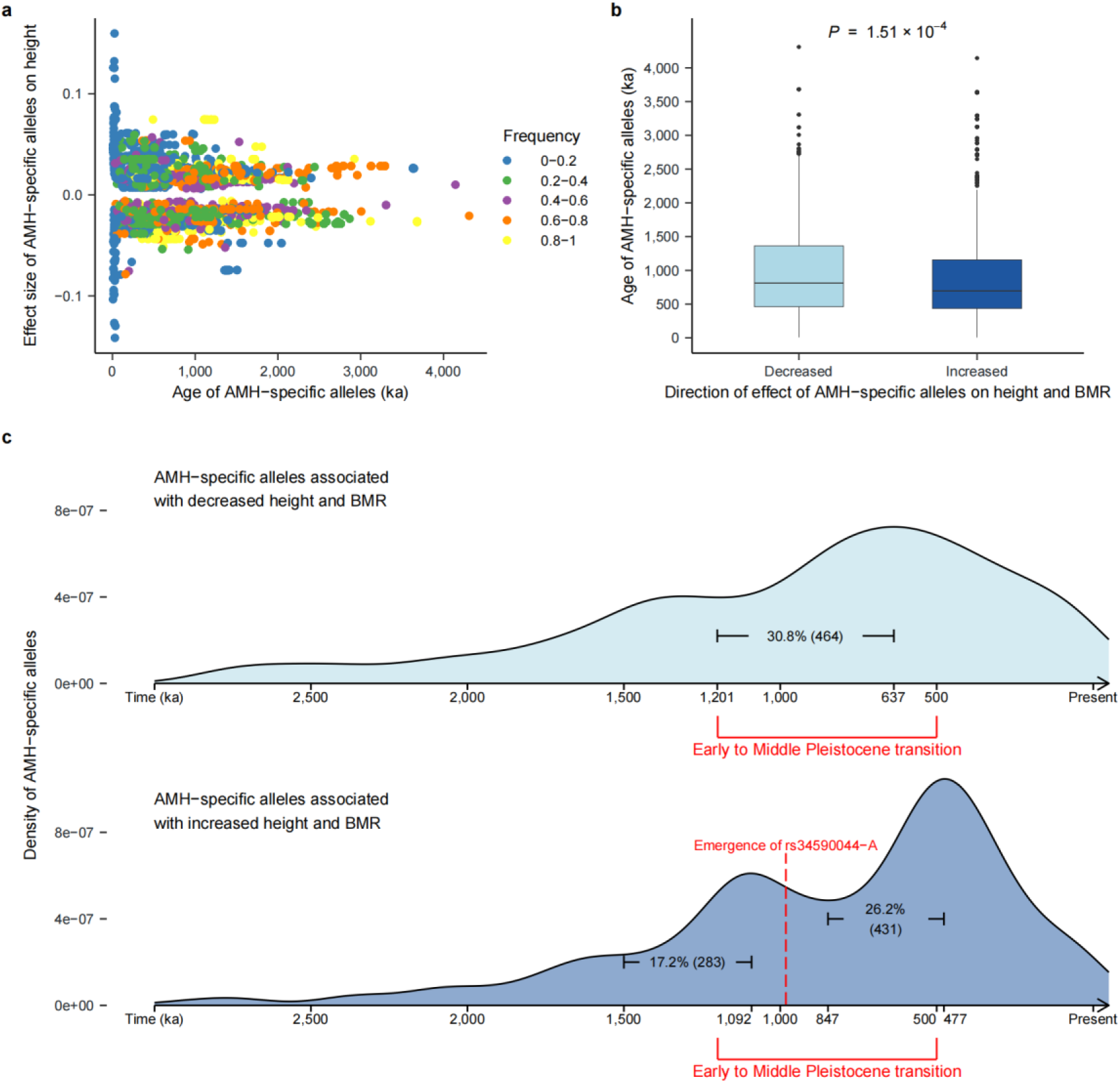
The Early to Middle Pleistocene transition has a strong influence to the evolution of inferred causal SNP for height and BMR. **a**: Comparison between effect sizes on height and the frequencies of the AMH-specific alleles of 3,150 SNPs in the present-day British (GBR) population. **b**: Comparison of ages of AMH-specific alleles (y-axis) stratified by the direction of effects on height and BMR (x-axis) in the GBR. A one-tailed Wilcoxon rank sum test was performed to compare the ages of AMH-specific alleles associated with decreased height and BMR to those associated with increased height and BMR, with the alternative hypothesis specifying “greater”. The raw *P* value is shown on top of the boxplot and the false discovery rate (FDR)-corrected *P* value is shown in **Supplementary Table 7**. The comparisons of other populations in the 1000 Genomes Project are shown in **Extended Data Fig. 1 and Supplementary Table 7**. **c**: Density of AMH-specific alleles with different ages in the GBR population. Only alleles that originated from three million years ago to the present (x-axes) are shown. Density of AMH-specific alleles (y-axes) associated with decreased (upper panel) and increased (lower panel) height and BMR are shown separately. The red brackets below the x-axes show the Early to Middle Pleistocene transition period. Three bursts of AMH-specific alleles and the corresponding number and fraction are shown inside each plot, with the time indicated on the x-axes.

Notably, we discovered a dynamic evolutionary trajectory for height and BMR, marked by three bursts of associated genetic mutations at different evolutionary time points (**Fig. 2c**). Two of these bursts occurred during the MPT. The most ancient burst commenced around 1.5 million years ago, peaking approximately 1 million years ago, during which an estimated 17.2% of the height and BMR-increasing mutations observed in the GBR population emerged. This was followed by a subsequent burst, starting around 1.2 million years ago and peaking at ∼635 kya, which accounted for 30.8% of the mutations associated with decreases in both height and BMR. The third and most recent wave began at 847 kya and peaked after the MPT, at around 473 kya, and was characterized by a burst of 26.2% of mutations linked to increased height and BMR. Our results suggest a significant environmental impact on human height and BMR during evolution.

### An AMH-specific mutation upregulates *ACSF3*

We explored the functional impacts of the 3,307 SNPs with AMH-specific alleles through a gene-based annotation using ANNOVAR^23^. In total, we identified 389 genes that were likely to be affected by 820 SNPs with AMH-specific alleles. Among these SNPs, 55 SNPs were nonsynonymous mutations in 52 genes, with one SNP leading to the loss of a start codon in *COQ10A*, and one SNP causing a gain of a stop codon effect in *DNAAF4* (**Supplementary Table 8**). In addition, 763 SNPs potentially regulate the expressions of 353 protein-coding genes based on the expression quantitative trait loci (eQTLs) annotation from the Genotype-Tissue Expression (GTEx) database^24^ and enhancer-gene-interaction annotation from the GeneHancer database^25^ (**Supplementary Table 8 and Methods**).

We then investigated the potential functional impacts when perturbing these 389 genes that are either harboring or regulated by the SNPs with AMH-specific alleles based on the information in the Mouse Genome Database^26^. We observed that the perturbations of 39 genes caused height-related (such as reduced body size or body length, abnormal axial skeleton morphology, and abnormal limb development) or BMR-related (such as increased basal metabolism, decreased respiratory quotient energy and decreased oxygen consumption) phenotypes in mice (**Extended Data Fig. 2a, Supplementary Table 9 and Methods**). However, we also observed that 86 genes have not been reported to be associated with either height- or BMR-related phenotypes in previous mouse knockout (KO) experiments^26^ (**Extended Data Fig. 2a and Supplementary Table 9**).

Among these genes, we found that *ACSF3* is particularly of interest for the following reasons. First, it is involved in the metabolism of meat-enriched macronutrients, including EAAs, odd-chain fatty acids, and cholesterol^12^, which is the key difference in diet between AMHs and other non-human apes^8^. Second, the elevated expression of *ACSF3* was observed in the liver of modern humans compared to other apes, including chimpanzees, bonobos, gorillas, and orangutans in prior studies^13,14^ (**Fig. 3a, Extended Data Fig. 2b and Methods**). Third, ACSF3 is a metabolic enzyme localized in the mitochondria, the center of cellular metabolism. Although the depletion of *ACSF3* leads to impaired respiratory rates^12^ and some variants in the coding region of *ACSF3* have been associated with height in humans with rare diseases such as CMAMMA^10^, most of them lead to decrease in expression of *ACSF3*^27^. In contrast, the causal variant(s) leads to elevated *ACSF3* expression observed in AMHs compared to other non-human apes, and its potential influences in the evolution of AMHs remain largely unexplored.

**Fig 3.**
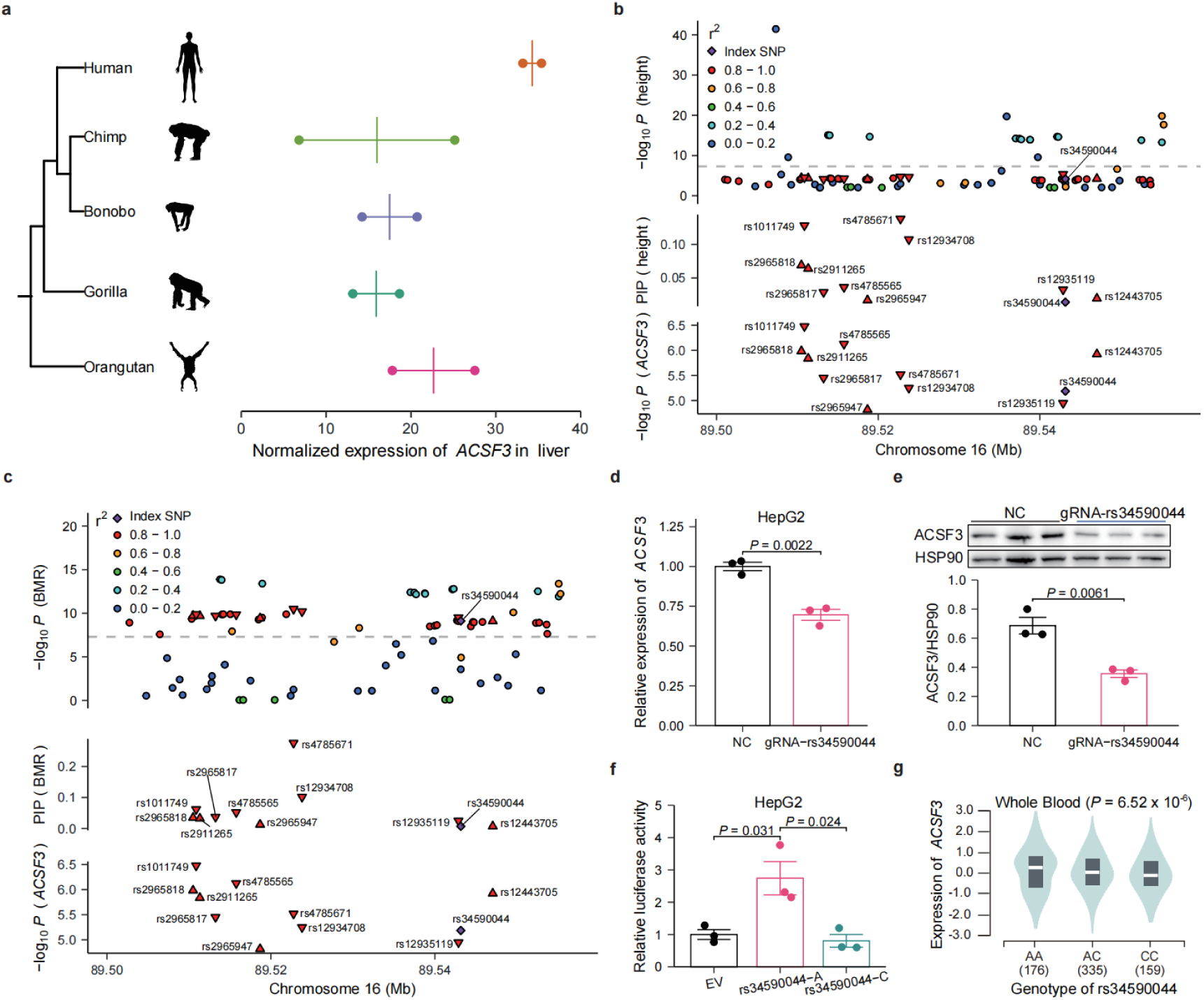
An AMH-specific mutation leads to increased expression of ACSF3. **a**: Expression of *ACSF3* in the liver of great apes. The x-axis shows both the mean expression (vertical lines) and the expressions of each individual (dots; N=2 for each species)^13^. The phylogeny is shown on the left (**Methods**). **b-c**: GWAS associations (top panel), fine-mapping (middle panel) and eQTL associations with *ACSF3* (bottom panel) of the 11 fine-mapped SNPs within one credible set. For GWAS associations with height (**b**) and BMR (**c**), SNPs within the genomic region (chr16: 89500494-89557054, extended 10 kbp upstream and downstream of the 11 fine-mapped SNPs; GRCh37) are also shown. For fine-mapping, the posterior inclusion probability (PIP) of each SNP is shown. For eQTL associations, the -log_10_(*P*) in whole blood in GTEx is shown. For all the panels, the purple rhombus represents the index SNP (rs34590044) and the color represents the LD (r^2^) between other SNPs and index SNP. For the 11 SNPs within the credible set, upper triangles represent increased effects and lower triangles represent decreased effects of AMH-specific alleles. **d)**: Expression levels of *ACSF3* when the region containing rs34590044 was inhibited by CRISPRi in HepG2 cells. NC: control. N=3. Error bars are 1 s.e.m. **e**: Protein levels of ACSF3 when the region containing rs34590044 was inhibited by CRISPRi in HepG2 cells. NC: control. N=3. Error bars are 1 s.e.m. **f**: Activity of the luciferase reporter of EV rs34590044-A or -C in HepG2 cells. Two-tailed t-test was performed between EV and rs34590044-A or rs34590044-A and rs34590044-C. N=3. EV: empty vector. Error bars are 1 s.e.m. **g**: Association between rs34590044 and the expression of *ACSF3* in whole blood in GTEx. The x-axis shows the genotypes of rs34590044 and the number of individuals and the y-axis shows the normalized expression of *ACSF3*. The raw *P* value is shown on the top. The figure is obtained from GTEx (**URLs**).

Here, we identified 11 potential causal SNPs with AMH-specific alleles of height and BMR at the *ACSF3* locus, all of which were in the non-coding regions, located within one credible set (**Fig. 3b and 3c**). Notably, they were annotated as eQTLs for *ACSF3* in the GTEx database (**Fig. 3b and 3c**)^24^. Interestingly, five AMH-specific alleles which were associated with the increased expression of *ACSF3* were also associated with increases in height and BMR (**Fig. 3b and 3c**). Conversely, the remaining six AMH-specific alleles were associated with the decreased expression of *ACSF3* and reduced height and BMR.

All these 11 candidate causal SNPs exhibit high LD with R^2^ > 0.93 (**Extended Data Fig. 2c**), which leads to none of the 11 SNPs having a PIP greater than 0.3. We thus sought to utilize functional approaches to pinpoint the causal SNP(s). To this end, we performed a CRISPR interference (CRISPRi)-based experiment in the liver-derived HepG2 cell line. Specifically, a set of nine guide RNA (gRNA) sequences was designed to silence the regions containing these 11 SNPs (**Supplementary Table 10**). While the gRNAs targeting other regions had no significant effects on the expression of *ACSF3* (**Extended Data Fig. 2d**), we observed that the gRNA targeting region chr16:89476746-89476963 (GRCh38) resulted in significant decreases in *ACSF3* expression (two-tailed *t*-test, *P* = 0.0023, **Fig. 3d**) and protein levels (two-tailed *t*-test, *P* = 0.0061; **Fig. 3e**) compared to the control, indicating that the SNP rs34590044 within this region is likely to be the causal SNP. In addition, the gRNA targeting rs34590044 had no significant effects on the expression of the other six genes in the proximal regions of the genome based on RT-qPCR experiments (**Extended Data Fig. 2e**). Furthermore, the AMH-specific allele, rs34590044-A, is significantly associated with increased enhancer activity compared to the ancestral rs34590044-C allele based on the luciferase experiment (**Fig. 3f**). This finding was corroborated by the GTEx^24^ and eQTLGen^28^ databases, wherein rs34590044-A was significantly associated with increased *ACSF3* expression (effect size = 0.103, *P* = 6.5 × 10^-6^ in GTEx; Z score = 4.3453, *P* = 1.4 × 10^-5^ in eQTLGen) in the whole blood (**Fig. 3g**). Together, our results suggest that the AMH-specific rs34590044-A allele can upregulate the expression of *ACSF3* compared to the non-AMH primates C allele.

### ACSF3 regulates mitochondrial activity

After confirming that rs34590044-A regulates the expression of ACSF3 by influencing its enhancer activity, we explored its functional impacts by perturbing ACSF3 expression in cellular models. The RNA-seq analysis revealed that the inhibition of ACSF3 transcription led to a dramatic change in the transcriptome (**Extended Data Fig. 3a-b**) characterized by a significant downregulation of genes, such as *TOMM22*, *NDUFB8*, *TIMM23*, *MTX1*, *CHCHD10*, *GEFR*, *FXN*, and *COX17*, related to the mitochondrial functions in the HepG2 cells (**Extended Data Fig. 3c-d**). In addition, we observed a significant reduction in the respiratory rate (two-tailed t-test, P = 0.0001), as the marker for mitochondrial activity^29,30^, of HepG2 cells subjected to ACSF3 expression inhibition using CRISPRi (**Extended Data Fig. 3e**). Thus, our results indicate that rs34590044-A acts as a key regulator for mitochondrial activity by regulating the expression of ACSF3.

We then investigated the potential mechanism by which ACSF3 influences the mitochondrial activity. As ACSF3 is responsible for the metabolism of macronutrients, we analyzed changes in cellular metabolism in HepG2 cells with targeted metabolomics. We found that the inhibition by CRISPRi induced a significant elevation in MMA (**Extended Data Fig. 3f and Supplementary Table 11**; two-tailed *t*-test, *P* < 0.0001), a cytotoxic metabolite. However, we did not observe significant changes in malonate, malonyl-CoA nor lysine malonylation (**Extended Data Fig. 3g-h**) when the expression of *ACSF3* was suppressed in the HepG2 cells. These results were different from the findings reported in a prior study using HEK293T derived from human embryonic kidneys^12^.

We then investigated the mechanism by which ACSF3 regulates mitochondrial activity *ex vivo* and *in vivo*. Via adenovirus infection, we identified that overexpression of *Acsf3* downregulated the abundance of MMA and enhanced the respiratory rates of the primary hepatocytes (**Fig. 4a-b, Extended Data Fig. 4a-b and Supplementary Table 12**). We then generated a *Acsf3* KO mouse model using CRISPR/Cas9 technology to induce mutagenesis in mice. As rs34590044 is located within a non-conserved region of the mouse genome (**Extended Data Fig. 4c**), we generated a whole-body depletion mouse model of *Acsf3* by depleting its second through fifth exons (**Fig. 4c**). The depletion of *Acsf3* in the liver (two-tailed *t*-test, *P* < 0.001 for RT-qPCR; two-tailed *t*-test, *P* < 0.0001 for western blot), brown adipose tissue (two-tailed *t*-test, *P* <0.001 for RT-qPCR; two-tailed *t*-test, *P* < 0.0001 for western blot), and muscle (two-tailed *t*-test, *P* < 0.001 for RT-qPCR, two-tailed *t*-test *P* < 0.0001 for western blot) was confirmed through RT-qPCR and western blot assays (**Extended Data Fig. 4d-e**).

**Fig 4.**
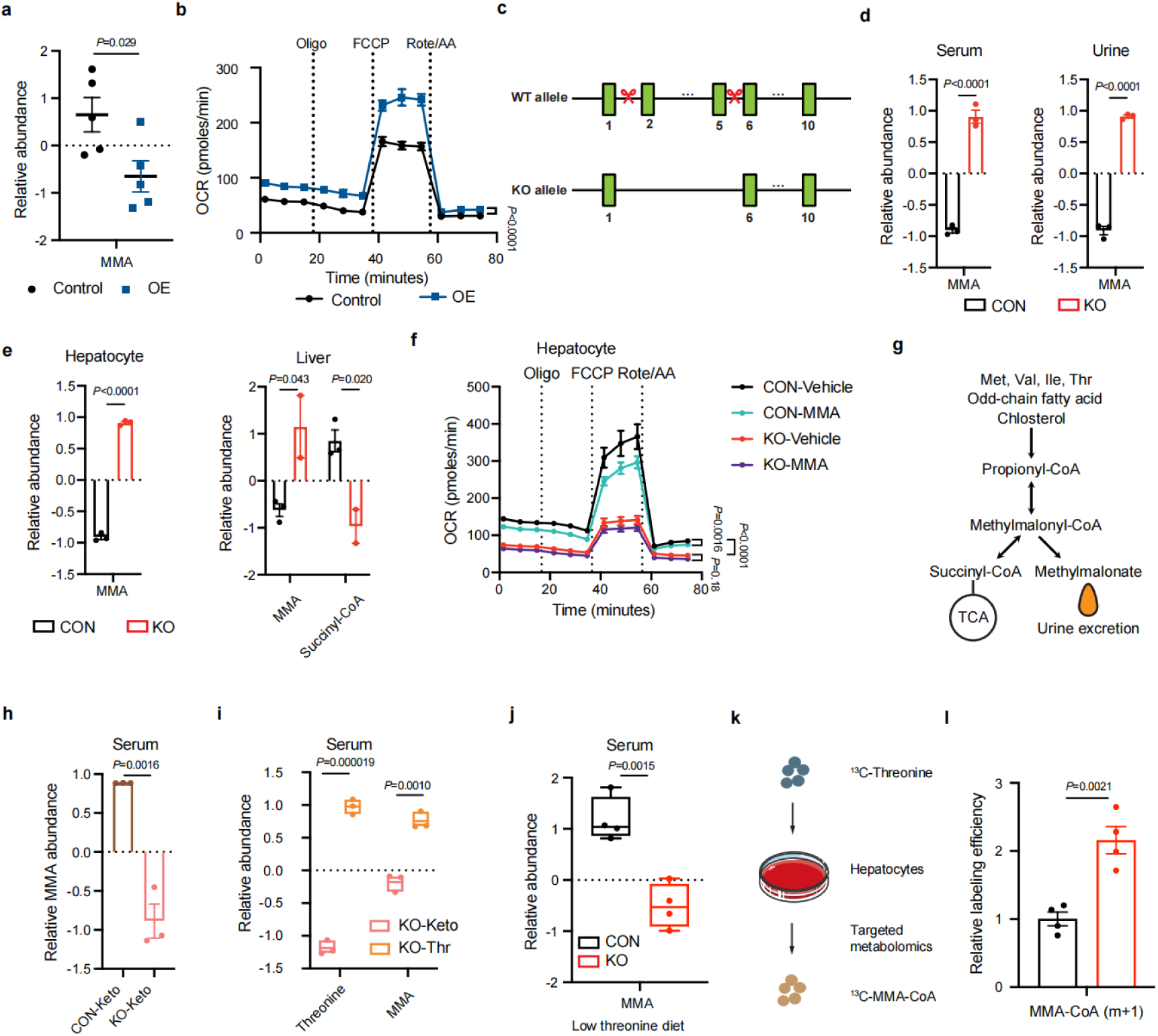
ACSF3 regulates mitochondrial activities via controlling the catabolism of threonine to MMA. **a-b**: Relative abundance of methylmalonate and respiratory rates in hepatocytes. CON: control. OE: adenovirus based *Acsf3* overexpression. MMA: methylmalonate. N=5. **c:** Establishment of the *Acsf3^-/-^* mouse model using CRISPR-Cas9 technology, in which exons 2 to 5 were depleted. **d:** Relative abundance of methylmalonate in serum and urine. CON: control mice (*Acsf3^+/-^*). KO: knockout mice (*Acsf3^-/-^*). MMA: methylmalonate. N=3. **e:** Relative abundance of methylmalonate in hepatocytes and methylmalonate+succinyl-CoA in the liver . CON: control mice (*Acsf3^+/-^*). KO: knockout mice (*Acsf3^-/-^*). MMA: methylmalonate. N=2–3. **f: R**espiratory rates in hepatocytes. MMA: 5 mM methylmalonate. N=3. CON: control mice (*Acsf3^+/-^*). KO: knockout mice (*Acsf3^-/-^*). N=2–3. **g: P**otential metabolic pathway of methylmalonate based on the literature^10^. **h:** Relative abundance of methylmalonate in the serum of mice administered a ketogenic diet. CON: control mice (*Acsf3^+/-^*). KO: knockout mice (*Acsf3^-/-^*). Keto: ketogenic diet. MMA: methylmalonate. N=3. **i:** Relative abundance of threonine and methylmalonate in mice administered a ketogenic diet and threonine in drinking water. KO: knockout mice (*Acsf3^-/-^*). Keto: ketogenic diet. MMA: methylmalonate.N=3. **j:** Relative abundance of methylmalonate in the serum of mice administered a low-threonine diet. CON: control mice (*Acsf3^+/-^*). KO: knockout mice (*Acsf3^-/-^*). MMA: methylmalonate. N=4. **k:** Scheme of a ^13^C-labeled threonine-based metabolic flux assay. **l:** Relative abundance of ^13^C-labeled methylmalonyl-CoA in hepatocytes. CON: control mice (*Acsf3^+/-^*). KO: knockout mice (*Acsf3^-/-^*). MMA-CoA: methylmalonyl-CoA. N=4.

Consistent with the observation from the immortalized HepG2 cells, the depletion of *Acsf3* in mice led to the significant elevation of MMA in the serum (two-tailed *t*-test, *P* < 0.0001), urine (two-tailed *t*-test, *P* < 0.0001), liver (two-tailed *t*-test, *P* = 0.043), and primary hepatocytes (two-tailed *t*-test, *P* < 0.0001) and the downregulation of succinyl-CoA in the liver (two-tailed *t*-test, *P* = 0.020) (**Fig. 4d-e and Supplementary Table 13-16**). In contrast, the depletion of *Acsf3* had no effects on the levels of malonate, malonyl-CoA or lysine malonylation in various types of tissues (**Extended Data Fig. 4f-g**). On one hand, the depletion of *Acsf3* significantly reduced the respiratory rate of the primary hepatocytes (**Extended Data Fig. 4h**). On the other hand, the *ex-vivo* MMA treatment remarkably decreased the respiratory rate of primary hepatocytes from wild-type but not *Acsf3^-/-^* mice (two-tailed *t*-test, *P* < 0.0001) (**Fig. 4f**). These results suggest that ACSF3 controls mitochondrial activities via regulating the metabolism of MMA.

We further understood the biochemical details regarding how ACSF3 regulates the metabolism of macronutrients. It has been reported that the catabolism of EAAs, cholesterol, or odd-chain fatty acid may contribute to the synthesis of MMA ^12^ (**Fig. 4g**). We fed the mice a ketogenic diet composed of 95% lipid and an extremely low amount of EAAs compared to the chow diet^31^. We observed that the feeding of a ketogenic diet diminished the elevation of MMA in the serum of *Acsf3^-/-^* mice (two-tailed *t*-test, *P* = 0.0016) (**Fig. 4h and Supplementary Table 17**), indicating that the ACSF3-dependent accumulation of MMA is associated with amino acid metabolism. Furthermore, the abundance of threonine, but not other EAAs, was significantly lower (two-tailed *t*-test, *P* = 0.0037) in the hepatocytes of *Acsf3^-/-^* mice compared to control mice fed a ketogenic diet (**Extended Data Fig. 4i**). The dietary supplementation of threonine via drinking water upregulated the serum level of MMA in the *Acsf3^-/-^* mice (two-tailed *t*-test, *P* = 0.001), but not in control mice, fed a ketogenic diet (two-tailed *t*-test, *P* = 0.29) (**Fig. 4i, Extended Data Fig. 4j and Supplementary Table 18**). In contrast, the dietary restriction of threonine diminished the elevation of MMA in the serum of *Acsf3^-/-^* mice (two-tailed *t*-test, *P* = 0.0016) (**Fig. 4j and Supplementary Table 19**). With a ^13^C-labeled threonine-based metabolic flux assay (**Fig. 4k**), we confirmed that the uptake of threonine (**Extended Data Fig. 4k**) and conversion rate from threonine to MMA was significantly accelerated by the depletion of *Acsf3* in the primary hepatocytes (two-tailed *t*-test, *P* = 0.0021) (**Fig. 4l and Supplementary Table 20**). These results demonstrated that ACSF3 regulated MMA metabolism via its effects on EAA catabolism, especially threonine.

### ACSF3 regulates body length and energy metabolism

Finally, we investigated the impacts of *Acsf3* on height (body length for mice) and BMR (energy expenditure for mice) *in vivo*. The *Acsf3^-/-^* mice exhibited normal viability and typical developmental progression. At three weeks of age, the body length of *Acsf3^-/-^* mice was similar to that of *Acsf3^+/-^* control mice (*Acsf3^+/-^*: 7.875 cm and *Acsf3^-/-^*: 7.850 cm). However, during the fourth through sixth weeks, *Acsf3^-/-^* mice exhibited slower growth, resulting in a significantly shorter body length (two-tailed *t*-test, *P* = 0.00072) compared to their littermate controls (*Acsf3^+/-^*) fed a normal chow diet (two-tailed *t*-test, *P* = 0.97, *P* = 0.14, *P* = 0.015, *P* = 0.0052) (**Fig. 5a-b**), without the obvious differences in body weight (**Extended Data Fig. 5a**). However, the differences in body length were diminished by feeding the mice a threonine restriction diet (two-tailed *t*-test, *P* = 0.040, *P* = 0.44, *P* = 0.28, *P* = 0.24) (**Fig. 5c-d and Methods**), demonstrating the significance of nutrients for body length. We also observed that the depletion of *Acsf3* downregulated energy expenditure (two-tailed *t*-test, *P* = 0.026) (**Fig. 5e**) without changes in the physical activities (**Extended Data Fig. 5b**) of mice fed a chow diet, as measured by a comprehensive laboratory animal metabolic system (CLAMS) assay.

**Fig 5.**
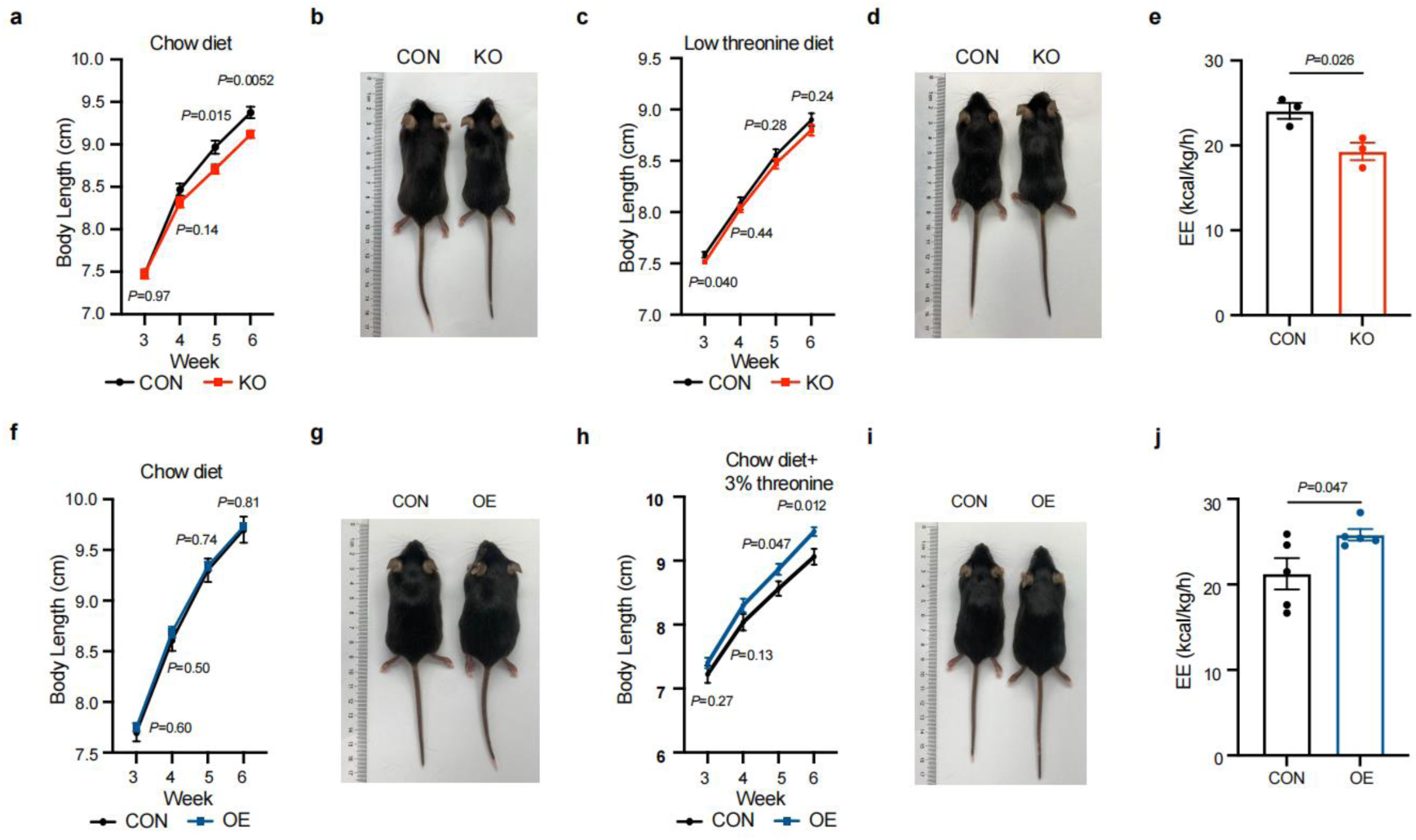
ACSF3 controls body length and energy metabolism in mice. **a-b**: Growth curve (**a**) and representative images (**b**) of body length in male mice administered a normal chow diet. CON: control mice (*Acsf3^+/-^*). KO: knockout mice (*Acsf3^-/-^*). N_CON_=12. N_KO_=11. **c-d:** Growth curve (**c**) and representative images (**d**) of body length in male mice administered a low threonine diet. CON: control mice (*Acsf3^+/-^*). KO: knockout mice (*Acsf3^-/-^*). N_CON_=13. N_KO_=18. **e:** Energy expenditure in male mice administered a normal chow diet. CON: control mice (*Acsf3^+/-^*). KO: knockout mice (*Acsf3^-/-^*). N=3. **f-g:** Growth curve (**f**) and representative images (**g**) of body length in male mice administered a normal chow diet. CON: injection of adeno-associated virus containing *GFP*. OE: injection of adeno-associated virus containing *Acsf3*. N_CON_=9. N_OE_=10. **h-i:** Growth curve (**h**) and representative images (**i**) of body length in male mice administered a normal chow diet and the dietary supplementation of threonine via drinking water. CON: injection of adeno-associated virus containing *GFP*. OE: injection of adeno-associated virus containing *Acsf3*. N_CON_=8. N_OE_=9. **j:** Energy expenditure in male mice administered a normal chow diet and the dietary supplementation of threonine via drinking water. CON: injection of adeno-associated virus containing *GFP*. OE: injection of adeno-associated virus containing *Acsf3*. N=5.

Similar phenotype characterization was performed on mice with the overexpression of *Acsf3*, which was initiated in the three-week-old mice. When the mice were fed a normal chow diet, the mice with *Acsf3* overexpression indeed presented similar body length and energy expenditure rate compared to the control mice (two-tailed *t*-test, *P* = 0.60, *P* = 0.50, *P* = 0.74, *P* = 0.81) (**Fig. 5f-g**). However, with the dietary supplementation of threonine via drinking water, the *Acsf3* overexpression mice presented longer body length (two-tailed *t*-test, *P* = 0.27, *P* = 0.13, *P* = 0.047, *P* = 0.012) (**Fig. 5h-i**) and higher energy expenditure rate (two-tailed *t*-test, *P* = 0.047) (**Fig. 5j**) with the dietary supplementation of threonine compared to control mice. The overexpression had no significant effects on the body weight or physical activities of mice (**Extended Data Fig. 5c-d**). These results indicate that ACSF3 controls the height and metabolism simultaneously, which has a close interplay with nutrition status.

### Distribution and evolution of rs34590044-A in humans

Next, we investigated the origin and geographic distribution of rs34590044-A in ancient and modern humans. We observed that the rs34590044-A is specific to AMHs and the ancestral allele, whereas rs34590044-C is conserved in the genomes of 19 non-AMH primates (**Fig. 6a**). Relate^21^ inferred the age of rs34590044-A at 981,916 years (with a lower age of 349,233 years and upper age of 1,614,601 years) in the GBR population, which coincided with the first burst of the variants associated with increased BMR and height (**Fig. 2b**). The ages of rs34590044-A range from 835,446 to 1,511,572 years in the other populations in the 1KGP (**Supplementary Table 21**), suggesting the rs34590044-A emerged after the divergence between humans and chimpanzees ∼6 million years ago but predated the split between AMHs and Neanderthals and Denisovans estimated at around 600 kya^32^. However, we observed that rs34590044-A is absent in the genomes of three high-coverage Neanderthal individuals^32–34^ and one high-coverage Denisovan^35^, as well as four additional low-coverage Neanderthal samples with a total of 16 reads (Goyet Q56-1, Mezmaiskaya 2, Les Cottés Z4-1514 and Denisova 11)^36,37^ (**Fig. 6b**). In contrast, rs34590044-A is common in the present-day GBR population, with an allele frequency of 54.4%^22^ as well as also at a high frequency across the other populations included in the 1KGP, with an average of 46.9% and a range from 21.2 to 65.3% (**Fig. 6c**)^22^.

**Fig 6.**
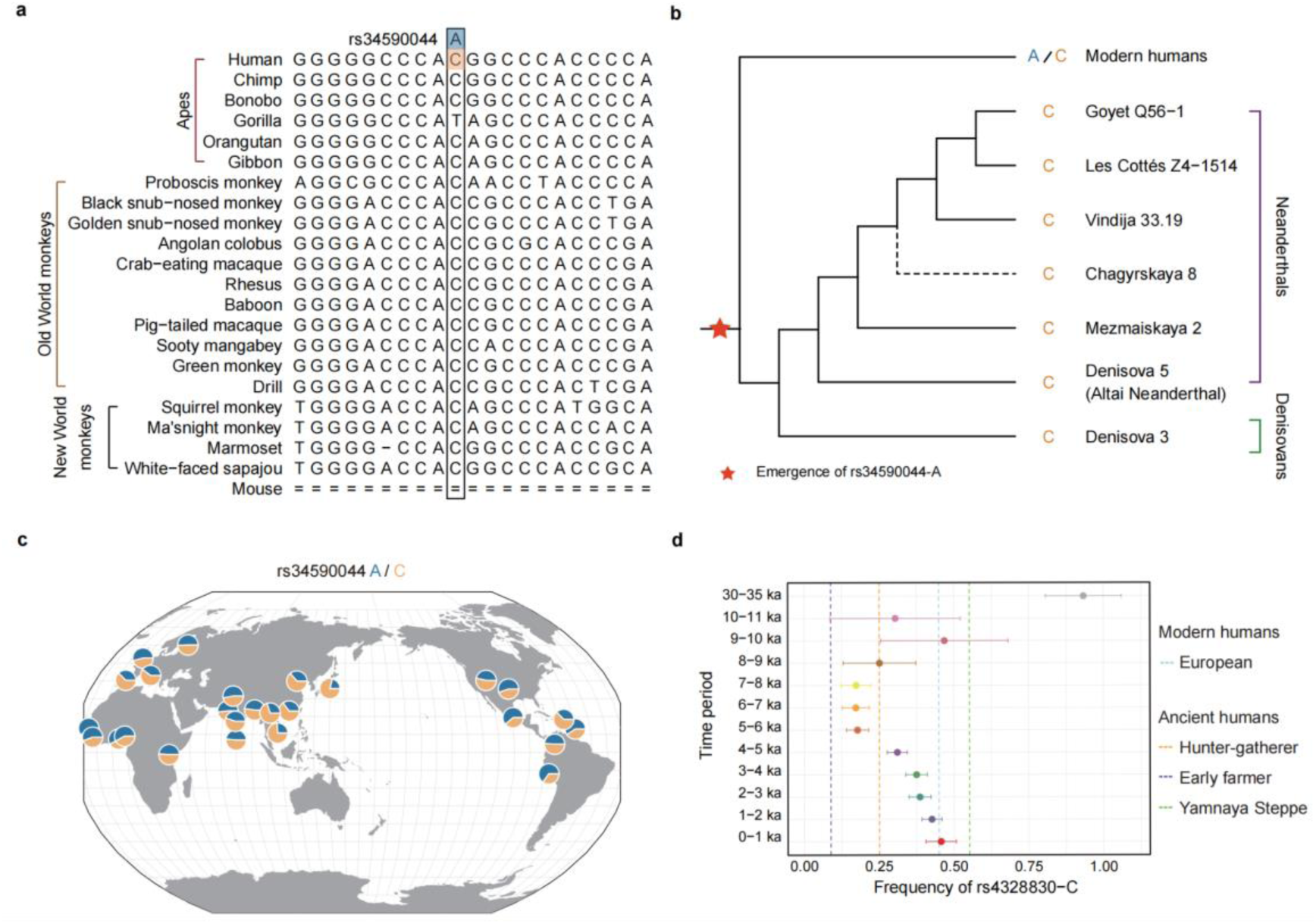
Distribution and evolution of rs34590044-A in modern and ancient humans. **a**: Alignment of rs34590044 in 21 primates and mouse in a 30-mammals Multiz Alignment (UCSC Genome Browser, GRCh38). The other six primates (tarsier, mouse lemur, sclater’s lemur, black lemur, coquerel’s sifaka and bushbaby) and two additional mammals (dog and armadillo) without aligned bases are not shown. **b**: Genotypes of rs34590044 in modern and archaic humans. Archaic humans include three high-coverage Neanderthal individuals (Vindija 33.19, Chagyrskaya 8 and Denisova 5 (the Neanderthal found in Denisova Cave at Altai Mountain)), three low-coverage Neanderthal individuals (Goyet Q56-1, Les Cottés Z4-1514 and Mezmaiskaya 2) and one high-coverage Denisovan. The lineage of Denisova 11 (with only the C allele) is not shown due to the mixture of Neanderthal and Denisovan ancestry^36^. The dashed line indicates the inferred lineage of Chagyrskaya 8 based on the split times between Chagyrskaya 8 and other Neanderthals^34^. The date where rs34590044-A originated is labeled with a red star. **c**: Allele frequencies of rs34590044 across 26 populations in the 1000 Genomes Project. Blue areas on pie charts indicate the AMH-specific derived A allele and orange areas on pie charts indicates the ancestral C allele. This figure was downloaded from the Geography of Genetic Variants Browser^61^. **d**: Allele frequencies of the rs4238830-C allele (in complete linkage with rs34590044-A) in ancient individuals in the past 35,000 years in Europe. The x- and y-axis show the frequencies and time periods, respectively. Dots represent the maximum likelihood estimates and error bars depict the 95% confidence intervals of the derived allele frequency. Vertical dashed lines indicate the frequency in modern European samples and the maximum likelihood estimates for ancient ancestries including the Hunter-gatherer group, Early farmer group and Yamnaya Steppe group.

We also studied the evolution of rs34590044 based on 6,474 ancient European human samples from the Allen Ancient DNA Resource (AADR) ^38^ (**Methods**). While rs34590044 was not included in the AADR dataset, we inferred its evolution history based on rs4238830, a SNP in complete LD with rs34590044 (D’=1, r^2^=1) in the GBR samples of the 1KGP.

We obtained the allele frequency distribution of rs4238830-C for ancient individuals across the past 35 ky (**Fig. 6d**). On average, rs4238830-C is at the highest frequency in the samples of Yamnaya Steppe (55%), followed by the European hunter-gatherers (25%), and Anatolian farmers (9%). The rs4238830-C allele was observed at a frequency of 85% in samples dating between 30 to 35 kya (**Fig. 6d**). We could not reliably infer the frequency of rs4238830-C from 20 to 30 kya due to less than 10 ancient samples being obtained at this interval. A dramatic decrease in the frequency of rs4238830-C was observed at 10 kya, declining from 85 to 25%, and the frequency was maintained at this level for the following two millennia. Between 8 and 5 kya, the frequency of rs4238830-C further decreased to 17%, potentially driven by the expansion of Anatolian Neolithic farmers into Europe^39^. Interestingly, from 5 kya until the present, the allele frequency exhibited a continuous increase reaching 45%, paralleling the frequency observed in present-day European populations. This trend corresponds with the population history of the Yamnaya steppe migration^40^ to Europe around the same period. Thus, our results suggested that the emergence and evolution of rs34590044 were shaped by both environmental and demographic factors.

## Discussion

In this study, we studied the genetic basis of height and BMR from an evolutionary perspective. Previous studies have revealed that these two traits have undergone intensive modifications, as modern humans possess a BMR three times higher than that of non-human apes and a general trend towards increased stature during the evolution of *Homo*. Although numerous genetic variants associated with adult height have been identified, these variants often cluster near skeletal growth genes^41,42^. Our study revealed a significant genetic correlation between height and BMR (**Fig. 1a and 1b**), indicating the existence of genetic factors with strong and simultaneous influences on both energy metabolism and height.

The three distinct bursts of height-BMR associated genetic variants, with two occurring during the Early to Middle Pleistocene transition (c. 1.2–0.5 Ma) and one following this critical transition period, indicated the significant impacts of the MPT on the evolution of human height and BMR (**Fig. 2b**). Previous research documented dramatic climate changes during the MPT, including a shift to longer glaciation cycles (from 41,000 to 100,000 years), more extreme thermal conditions, colder sea surface temperatures, and severe drought^43^, which profoundly influenced the distribution and evolution of biota, including the ancestors of modern humans^43^. For example, a severe population bottleneck was observed from 930,000 to 813,000 years ago, that reduced the effective population size of human ancestors from about 100,000 to only around 1,280 breeding individuals^44^. This bottleneck is also likely to lead to a gap in the availability of African and Eurasian fossil records. Notably, we found that a burst of negative variants accounting for 30.8% of the causal variants negatively associated with height and BMR that emerged during the 1.2 million to ∼635 kya period, coinciding with the MPT. Individuals harboring these variants conferring lower height and BMR would have had reduced food requirements, potentially enhancing their chances of survival during this challenging climatic transition.

Immediately after the MPT, we observed a burst of variants associated with increased height and BMR within a period of 300 ky from 700-400 kya. The stable climate during this period could have facilitated a more consistent food supply, rendering the higher energy requirements associated with increased BMR and taller stature less of a disadvantage. Additionally, a higher BMR allows increased energy expenditure, which may have conferred advantages for hunter-gatherer populations by improving their mobility and endurance.

Our results based on cellular and mouse models identified a non-coding AMH-specific mutation rs34590044-A, which coincided with the first burst of variants, contributing to the changes in human height and BMR. We observed that rs34590044-A is associated with the increased enhancer activity and expression of *ACSF3* compared to the ancestral allele conserved in the genomes of 19 non-AMH primates (**Fig. 3**). Our *in vitro*, *ex vivo*, and *in vivo* experiments demonstrated that the elevated expression of ACSF3 could contribute to increased body length and energy expenditure in a threonine-dependent manner (**Figs. 4-5**). Meat-enriched diets have ample EAAs including threonine. We observed that the higher expression of *ACSF3* in the AMHs could catalyze threonine into the TCA cycle, a key energy resource in living cells. In addition, elevation of *ACSF3* expression can avoid the accumulation of MMA, a toxic metabolite impairing mitochondrial activities. Therefore, rs34590044-A is likely to contribute to the transition from an herbivorous to a carnivorous diet in AMHs.

The degree and timing of the transition from an herbivorous ape-like diet to a more omnivorous diet rich in animal protein remains a subject of debate. Nevertheless, archaeological and anthropological evidence suggests that this shift occurred gradually over an extended period during human evolution, potentially dating back more than 3 million years ago^8,45^. The transition was thought to be induced by the expansion of drier grasslands and semi-forested regions, where digestible plant foods became less readily available compared to wetland forests, but grazing animals were abundant. In contrast, the inferred age of the rs34590044-A allele is approximately one million years, implying a prolonged process of genetic adaptation an omnivorous diet throughout human evolution.

Although the rs34590044-A mutation emerged before the divergence of modern humans, Neanderthals, and Denisovans from their common ancestor, we observed that it is absent in the genomes of Neanderthal and Denisovan individuals (**Fig. 6**). There are several potential explanations for this observation. First, the ancestors of Neanderthals and Denisovans may not have carried the rs34590044-A allele when they diverged from the common ancestor shared with AMHs. Alternatively, the rs34590044-A variant may have been present in the gene pools of Neanderthal and Denisovan ancestral populations but was subsequently eliminated due to genetic drift or selective pressure against this mutation. A third possibility is that rs34590044-A is present in the genomes of Neanderthals and Denisovans but has not yet been detected due to the limited number of ancient genomic samples currently available for these archaic human lineages. Further investigation of a more comprehensive archaic DNA dataset may help to elucidate the precise evolutionary dynamics and distribution of rs34590044-A across modern and archaic human populations.

Consistent with our hypothesis that the rs34590044-A allele may contribute to the human adaptation to a meat-rich diet, we observed that the frequency of this allele aligned well with the level of meat consumption in ancient European populations. The allele exhibited substantially higher frequency in the Yamnaya Steppe population (55%) and European hunter-gatherers (25%), both of whom likely subsisted on diets rich in animal-based proteins, compared to the Anatolian farmers (9%), whose diet was predominantly plant-based. Furthermore, our results suggested that demographic factors played a crucial role in shaping the frequency of the rs34590044-A allele and its associated physiological traits over the past 30,000 years in these ancient European populations (**Fig. 6**).

In summary, our findings of the present study unveiled the roles of complex evolutionary processes and the intricate interplay between genetic, environmental, and demographic factors in shaping crucial phenotypes and physiological traits within AMHs. The results highlight the multifaceted nature of human adaptation and underscore the importance of considering diverse influences when examining the origins and prevalence of key physiological characteristics.

## Methods

### GWAS summary statistics

We obtained GWAS summary statistics of height and BMR from Loh et al^9^ (URLs), which are produced by BOLT-LMM with a linear mixed model using up to 459,327 samples from UK Biobank. There are no known limitations or data transfer agreements regarding access to the summary data described above. The GWAS summary statistics were then filtered to minor allele frequency (MAF) >= 0.01 and INFO score (imputation quality) >= 0.9 for our further analyses.

### Estimation of global genetic correlation

We first used the munge_sumstats.py of LDSC v1.0.1^15^ with default parameters to munge the raw GWAS summary statistics. In total, 6,319,071 and 6,319,157 SNPs for height and BMR are passed through the filtering criteria, respectively.

We then used the ldsc.py script of LDSC v1.0.1^15^ to calculate the global genetic correlation between height and BMR from the munged summary statistics. Specifically, we used the following command:

python -u ldsc.py \
--rg "${path_to_munged_sumstats}" \
--ref-ld-chr "${eur_ref_ld_chr}" \
--w-ld-chr "${eur_ref_ld_chr}" \
--out "${out_prefix}"
where ${eur_ref_ld_chr} referred to 1000G_Phase3_ldscores.tgz, which was downloaded from Gazal et al ^46^ (URLs).

### Estimation of local genetic correlations

We estimated the local genetic correlations using LAVA v0.1.0^17^ across 2,495 genomic loci (URLs). The genomic loci were created by LAVA to ensure the minimal LD between them while keeping approximately equal size. As the authors of LAVA suggested, we used LDSC to estimate sample overlap between summary statistics of height and BMR. Custom scripts were used (URLs) to run LAVA. In brief, univariate tests were performed to filter out loci lacking significant genetic signals (i.e., local heritability) for either trait. Then the loci which passed through Bonferroni corrections of univariate tests were used to calculate the local genetic correlations.

### Statistical fine-mapping of loci with significant local genetic correlations

We utilized SuSiE v0.12.40^18^ to perform statistical fine-mapping at loci with significant local genetic correlations between height and BMR. SuSiE requires both GWAS summary statistics and LD information (either from the same sample as GWAS, i.e. in-sample LD, or from a suitable reference panel^47^) to estimate potential causal variants. We used the in-sample LD estimated from 337,491 UKBB samples by Weissbrod et al^19^.

To run SuSiE, we used a custom pipeline from a prior study^48^, which infers the causal SNP within each locus by using the susie_rss function. We ran this pipeline using default default values for all parameters except for the L and n-samples parameters. For the L parameter, which represents the maximum number of causal SNPs at each locus, we specified 10 as previously suggested ^19,48^. SuSiE will estimate the true number of causal SNPs ranging from one to L. For the n-samples parameter, we used the maximum number of samples of GWAS of each trait (N = 458,303 for height and 451,316 for BMR, respectively).

SuSiE will report the 95% credible set (CS) at each locus, where each CS represents the minimum set of SNPs that contains the true causal SNP with a probability greater than 95%. Where multiple CSs exist at one locus, there are multiple causal SNPs. We then used SNPs within the 95% credible set (CS) from SuSiE as candidate causal variants in the downstream analyses.

### Identification of human-specific alleles

To facilitate comparisons between SNPs in humans with genomes of non-human primates (NHPs), we obtained the Multiz Alignment of 30 mammals (including human (GRCh38), 26 NHPs, mouse, dog and armadillo) from UCSC (URLs). We converted the genomic positions of shared candidate causal SNPs of height and BMR from GRCh37 to GRCh38 using the dbSNP155 database as UCSC suggested^49^. We then extracted the homologous bases of the 26 NHPs from Multiz Alignment for the shared candidate causal SNPs. SNPs that do not have aligned bases in NHPs were considered as human-specific insertions. For SNPs that have aligned bases in other NHPs, only bases that do not occur in any genomes of 26 NHPs are considered as human-specific mutations.

### Allele ages in the 1KGP by Relate

We obtained allele ages estimation for the shared candidate causal SNPs of height and BMR from Relate^21^ (URLs). We calculated the lower age and upper age as well as the age estimate for a neutral mutation as described previously^21^. To obtain years from generations, we multiply by 29^50^.

### Gene prioritization

For the identification of target genes of shared potential causal SNPs with human-specific alleles between height and BMR, we utilized three steps as below:

We first performed a gene-based annotation with RefSeq genes through ANNOVAR v2020-06-08^23^ to identify whether SNPs cause protein coding changes and the corresponding amino acids that are affected (i.e., missense SNPs that cause an amino acid change, lead to the immediate creation or loss of stop codon at the variant site or lead to the immediate creation or loss of start codon at the variant site);

For SNPs that cause protein coding changes, we refer to the affected genes as their target genes;

For SNPs that do not cause protein coding changes (synonymous SNPs within coding regions or SNPs within noncoding regions), we consider that these SNPs may possess regulatory potentials to affect the expression of nearby genes.

To identify target genes of SNPs that do not cause protein coding changes, we first collected cis-eQTLs in 49 tissues from GTEx v8^24^ and enhancer-gene interactions predicted by GeneHancer database^25^. Then we used a conservative method to locate the potential target genes. Specifically, we localized SNPs that are associated with expressions of at least one gene in at least one tissue (i.e., cis-eQTLs of genes) and that are located within enhancer regions. For SNPs that are cis-eQTLs, the genes whose expression is associated with SNPs are considered as the target genes of that SNPs. For SNPs located within enhancers, the predicted genes of enhancers are considered as the target genes of that SNPs. We subsequently constructed a SNP-to-gene map, ensuring the inclusion of the same target genes identified through both cis-eQTLs and enhancers associated with the SNPs.

### Classification of candidate genes based on MGI database

We utilized the Mouse Genome Informatics (MGI) database^26^ to investigate the functional impact of candidate genes.

We first collected the phenotypes and genotypes of mice with genes knocked out from the MGI database (HMD_HumanPhenotype.rpt and MGI_PhenoGenoMP.rpt; URLs). After removing eight genes that do not have homologs in mice and 90 genes that induce no observable phenotypes, we grouped the following 291 genes into five categories as below:

154 genes that lead to lethal effects (whose MGI terms contain "lethality" or "premature death") in mice;

12 genes that lead to infertile effects (whose MGI terms contain "infertility") in mice;

33 genes that caused height-related phenotypes (whose MGI terms contain one of the following keywords: "limb", "skeleton", "bone", "body size", "body length", "body height", "dwarf", "tibia", "femur", "fibula", "vertebra", "ilium") after perturbation in mice;

6 genes that caused BMR-related phenotypes (whose MGI terms contain one of the following keywords: "oxygen consumption", "respiratory quotient", "basal metabolism", "carbon dioxide production", "energy expenditure", "energy homeostasis", "energy metabolism", "adaptive thermogenesis") after perturbation in mice;

86 genes that have not been reported to be associated with either lethality, infertility, height or BMR-related phenotypes.

### Evolution of rs4238830 in ancient human samples

To investigate the evolution history of rs4238830 (as a proxy of rs34590044 based on complete LD between them), we collected ancient DNA data from AADR version v54.1 dataset^51^, and calculated allele frequency for 6474 ancient Europeans spanning from 150 BP (Before the Present) to 34948 BP. For allele frequency estimates, we grouped ancient individuals into groups per millennium, and kept groups with more than 10 individuals for downstream frequency calculation. In the time period with fewer than 10 ancient individuals, for instance, in each millennium between 30,000 BP to 35,000 BP, we grouped all available ancient individuals from Europe into one group for downstream calculation. The total number of individuals in each group is summarized in **Supplementary Table 22**.

Following equations and distributions as described in Mathieson et al^52^, we calculated allele frequency and related confidence intervals per group to examine the allele frequency distributions over time.

For reference, we collected the allele frequency information of modern populations from Europe calculated from 1000 Genomes Project Phase 3^22^, namely EUR. Given the genetic ancestry of modern Europeans could be traced back to three ancestral populations related to Early farmer, Yamnaya Steppe, and Hunter-gatherers, we calculated allele frequency for each ancestral European population for reference, grouping 61 ancient European individuals into group Early farmer, 75 individuals into group Yamnaya Steppe, and 64 individuals into Hunter-gatherers group (**Supplementary Table 23**).

### The silhouette of human and non-human primates

The silhouette of human is created by Katy Lawler (http://phylopic.org/; under the license of https://creativecommons.org/licenses/by/4.0/); the silhouette of chimpanzee is created by Kai R. Caspar (http://phylopic.org/; under the license of https://creativecommons.org/licenses/by/3.0/); silhouettes of bonobo, gorilla and Orangutan are from http://phylopic.org/ under a Public Domain Dedication 1.0 license.

### Reagents

**Supplementary Table 24** contains all reagents in the present study.

### Procedures involving animals

All the procedures involving animals were performed in accordance with the guidelines of, and approved by, the Fudan University ethical committee. Both sexes were included in the study. The *Acsf3* KO mice (C57 B6/J) were obtained from Gem Pharmatech (Nanjing, China). Mice were maintained in a SPF facility under a 12 h light/12 h dark cycle, at a specific pathogen free facility at constant temperature of 23°C, and with free access to food and water. The ketogenic diet was purchased from BioServe (S3666), and 3% threonine was administered via drinking water. The low threonine diet (A23092101) and corresponding control diet (A17100503) were customized from SYSE BIO. The specific ingredients and caloric value of these two diets were listed in **Supplementary Table 25**.

### RT-qPCR

Total RNA was extracted from tissues using RNAiso Plus (Takara). The extracted RNA was quantified by Nanodrop and 500 ng of it was converted into cDNA using a PrimeScript™ RT reagent Kit (Takara). Quantitative RT-PCR (qRT-PCR) was performed using SYBR Green PCR Master Mix (Takara) in an Applied Biosystems QuantStudio 3. *36b4* or *ACTB* was used as an internal reference gene. Primer sequences are provided in **Supplementary Table 26**.

### Western blotting

Tissues were lysed in 1% NP-40 buffer (50 mM Tris-HCl pH 7.5, 150 mM NaCl, 1% Nonidet P-40) supplemented with protease and phosphatase inhibitors (ApexBio Technology). Proteins were then separated using 10% SDS-PAGE and transferred to PVDF membranes (Merck Millipore). PVDF membranes were then blocked by 5% BSA, incubated with the various specific primary antibodies overnight at 4°C, and then with HRP-conjugated secondary antibodies for 60 min at room temperature (**Supplementary Table 27**). ECL Reagent (Yeasen) and Clinx ChemiScope 6000 (Clinx Science Instruments Co., Ltd.) were used for visualizing protein bands.

### Mitochondrial purification

Tissues were homogenized in 300 μL/50 mg ice-cold hypotonic buffer (10 mM KCl, 1.5 mM MgCl2, 10 mM Tris-HCl pH 7.4) in a glass homogenizer using an application of 30 strokes; debris were then removed by filtration. The suspension was centrifuged at 1,000 × g for 10 min at 4°C. The upper layer of the suspension was transferred into a clean centrifuge tube and centrifuged at 10,000 × g for 30 min at 4°C. The mitochondrial pellet was lysed in 500 μL 1% NP-40 buffer by gently mixing for 30 min at 4°C, and then centrifuged at 12,000 × g for 15 min at 4°C. The supernatant contained mitochondrial proteins.

### Isolation of primary hepatocytes

Primary hepatocytes were isolated from 6 to 8-week-old male mice. Briefly, the mice were anesthetized with 1% pentobarbital sodium (40 μL per 10 g body weight, delivered using intraperitoneal injection). Then, the mice were perfused with Krebs-Ringer Solution with Glucose Buffer (KRG Buffer: 120 mM NaCl, 24 mM NaHCO3, 20 mM glucose, 5 mM HEPES, 4.8 mM KCl, 1.2 mM MgSO4, 1.2 mM KH2PO4) containing 0.1 mM EGTA from the hepatic portal vein. The liver tissue was then digested with a KRG buffer containing 2 mM CaCl2 and 0.5 mg/mL collagenase type IV (Sigma-Aldrich). Then, the liver tissue was minced to release the hepatocytes^53^. Consequently, the hepatocytes were isolated by gradient density centrifugation using Percoll beads (GE HealthCare, USA) and pelleted by centrifugation (1000 × g, 2 min). The pellets were then resuspended in 1 g/L DMEM (Gibco) containing 10% FBS (Gibco), and 1% penicillin-streptomycin (Thermo Fisher), and plated at 400,000 cells/mL for further assays.

### Seahorse assay

Oxygen consumption was measured using a Seahorse XFp Analyzer (Agilent Technologies). First, 32,000 cells were plated in poly-L-lysine-coated XFp cell culture miniplates and incubated overnight. Concurrently, a sensor cartridge was hydrated overnight in a 37°C incubator without CO_2_. Cells were then incubated with assay media (XF base medium supplemented with 25 mM glucose, 2 mM L-glutamine, 1 mM pyruvate, and 5 mM HEPES) in an incubator at 37°C without CO_2_ for 1 h prior to conducting the assay. The A, B, and C ports of the cartridge were loaded as the following inhibitors were added sequentially at the specified concentrations: 10 μM oligomycin, 10 μM FCCP, and 5 μM rotenone/antimycin A. Oxygen consumption was measured repeatedly using the corresponding program according to the manufacturer’s protocol. Cell number was determined by Countess 3 (Invitrogen) and was used to normalize oxygen consumption rate in pmoles/min.

### CRISPRi

sgRNAs were designed using the CRISPick tool from the Broad Institute^54^. All sgRNA sequences are listed in **Supplementary Table 10**. HepG2 cells were first infected with a lenti-dCas9-KRAB-MeCP2-blast vector and selected using 10 µg/mL blasticidin for at least 7 days. HepG2 cells stably expressing dCas9-KRAB-MeCP2 were then infected with a LentiGuide-Puro vector carrying either non-targeting or SNP-targeting sgRNAs, and selected using 2 µg/mL puromycin for at least 2 days prior to further molecular or cellular assays.

### Luciferase reporter assay

Luciferase reporter assays were performed as described previously^55^. The enhancer luciferase reporter constructs with the control luciferase construct Renilla were co-transfected into cells using PEI (Polysciences). The luciferase signal was first normalized to that of the Renilla luciferase signal, and then to that of the empty pGL3 minimal promoter vector.

### Metabolite extraction

Extraction method was modified from that used in a prior study^56^. Briefly, for the serum, the mouse blood was centrifuged at 14,000 × g for 10 min at 4°C, and supernatant was decanted into a new centrifuge tube. Then, 80% methanol (H_2_O: MetOH = 1:4) was added into the centrifuge tube and vortexed for 1 min. For the cells, a total of 100 µL acetonitrile/water (50/50) was used to reconstitute the aqueous metabolites. The mixture was incubated for 30 min at −80°C, and then centrifuged at 4000 × g for 10 min at 4°C to separate the aqueous metabolites. Supernatant was collected and dried under nitrogen gas at room temperature. The dried metabolites were stored at −80^°^C until analysis using LC-MS.

### Targeted metabolomics

Aqueous metabolites were reconstituted using 100 µL acetonitrile: water (*v:v* 50:50). Then, 5 µL reconstituted sample was injected into the LC-MS. The method used was modified from a previously published protocol^57^ that uses an amide HILIC column (XBridge Amide 3.5 µm, 4.6 × 100 mm). Briefly, we used two elution solutions: buffer A (95% water and 5% acetonitrile with 20 mM ammonium hydroxide and 20 mM ammonium acetate, pH 9.0) and buffer B (acetonitrile). The 0.25 mL/min LC gradient was initiated from 0–0.1 min, 85% B, 3.5 min, 32% B, 12 min, 2% B, 16.5 min, 2% B, and 17–16 min, 85% B. Samples were acquired by QTRAP 5500+ (AB Sciex) using a polarity switching approach adapted from a published MRM list containing 297 transitions. LC-MS/MS peak integration was performed on a MultiQuant (AB Sciex) to obtain the relative abundance of metabolites via a spreadsheet.

### 13C-labeled glucose/lactate/glutamine treatment

The hepatocytes were seed in 10mm plates at a density of 400,000/mL. Fasting with high DMEM medium without serum for 12h, the cell culture media were replaced by complete DMEM containing ^13^C-labeled threonine (95 mg/L) and extracted cell metabolites. The analysis by targeted Metabolomics was performed on Sciex OS as **Supplementary Table 28**.

### RNA-seq Assay

Total RNA was extracted from HepG2 using TRIzol reagent (Thermo Fisher Sci-entific). Extracted RNA was then reversely transcribed using the PrimeScript™ RT reagent Kit (Takara). RNA-seq libraries for expression analysis were constructed using KAPA RNA Hy-perPrep Kit KR1350 v1.16 according to the vendor’s protocol and paired-end 2 x 150 bp reads were sequenced using the Illumina HiSeq platform. The raw data were aligned and quantified by HISAT2^58^. The raw data was deposited to Gene Expression Omnibus at GSE203040. Differentially expressed genes were determined by DESeq2^59^ and *P* value < 0.05 and Log2 fold change > 2 were recognized as the cutoff.

### Body Composition

Body composition was acquired using an Echo MRI (Echo Medical Systems, Houston, Texas) with a 3-in-1 Echo MRI Composition Analyzer in order to determine lean mass and fat mass values prior to biochemical experiments.

### Indirect calorimetry

The energy expenditure was determined by the Promethion Comprehensive Lab Animal Monitoring System (CLAMS, Sable Systems International, NV, USA) housed within a temperature-controlled environmental chamber at Fudan University. Before the experiment started, the mice were acclimatized in the CLAMS for 48 h and monitored for 48h. The data were processed as previously described^60^.

### Statistical analysis

We conducted all statistical analyses using R version 4.1.0.

### URLs

GWAS summary statistics of height and BMR from Loh et al are obtained from https://alkesgroup.broadinstitute.org/UKBB/; LD scores of samples with European ancestry in 1000 Genome Project Phase 3 are downloaded from https://zenodo.org/records/7768714; 2,495 genomic loci with minimal LD are obtained from https://github.com/cadeleeuw/lava-partitioning; Custom scripts to run LAVA is at https://surfdrive.surf.nl/files/index.php/s/rtmNm8YZERGwl7f?path=%2Fcluster_setup%2Fscripts; UCSC hg38 multiz30way alignments are downloaded from https://hgdownload.soe.ucsc.edu/goldenPath/hg38/multiz30way/maf/; Allele ages in the 1KGP estimated by Relate are obtained from https://zenodo.org/records/3234689. Phenotypes and genotypes of mice in MGI are downloaded from https://www.informatics.jax.org/downloads/reports/index.html.

## Acknowledgements

We thank Xia Shen at Fudan University, Yuanqing Feng at University of Pennsylvania and Jeffrey P. Spence at Stanford for commenting on the manuscript. We thank Single Cell Quantitative Metabolomics and Lipidomics Core Facility of IMIB at Fudan University for LC-MS/MS analysis. Funding: This work was supported by the following grants: MOST 2020YFA0803600, 2018YFA0801300, NSFC 32071138, and SKLGE-2118 to J.L.; MOST 2020YFA0803800, 2019YFA0801900, NSFC92057115, and the Shanghai Sailing Program 20YF1402600 to H.H.; grants from the National Key R&D Program of China (2020YFE0201600 and 2021YFC2500202), National Natural Science Foundation of China (grant No. 31970563), the 111 Project (B13016), and Shanghai Municipal Science and Technology (grant No. 2017SHZDZX01, grant No. 19410741100) to S.F.

## Author contributions

Conceptualization, Y.Z., J.W., C.Y., Y.S., Z.Y., S.Z., K.W., H.H., J.L. and S.F.; Investigation: Y.Z., J.W., C.Y., Y.S., Z.Y., S.Z., K.W., H.H., J.L. and S.F.; Writing, Y.Z., J.W., C.Y., Y.S., Z.Y., S.Z., K.W., H.H., J.L. and S.F.; Visualization, Y.Z., J.W., C.Y., Y.S., Z.Y., S.Z., K.W., H.H., J.L. and S.F.; Funding Acquisition, H.H., J.L. and S.F.; Supervision, H.H., J.L. and S.F..

## SUPPLEMENTARY MATERIALS

**Extended Data Fig 1.**
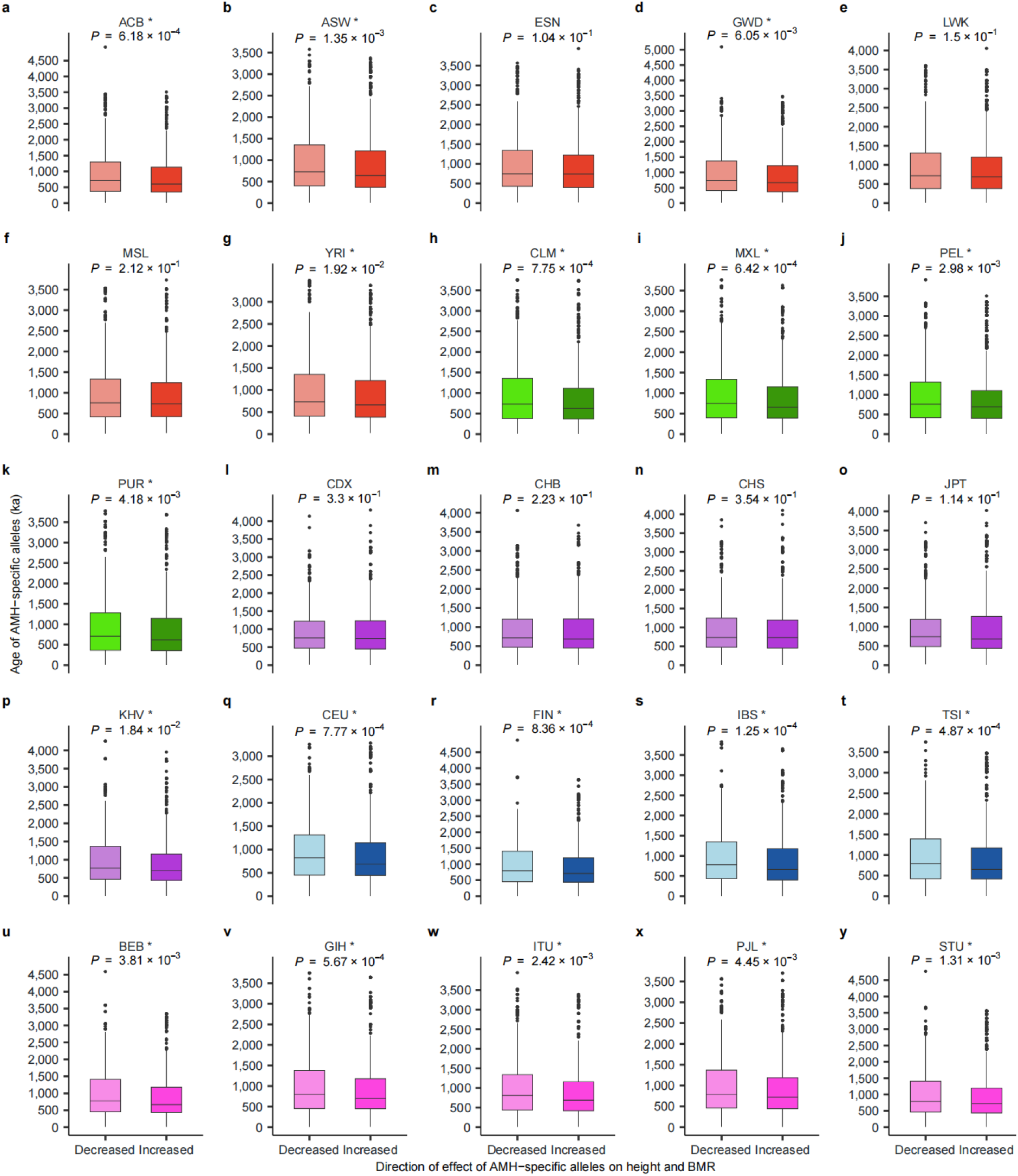
Comparison of ages of AMH-specific alleles (y-axis) stratified by the direction of effects on height and BMR (x-axis) in 25 populations in the 1KGP. One-tailed Wilcoxon rank sum test is performed to compare the ages of AMH-specific alleles associated with decreased height and BMR to those associated with increased height and BMR, with the alternative hypothesis specifying “greater”. Raw *P* values are shown on top of the boxplot and FDR-corrected *P* values are shown in **Supplementary Table 7**. Populations with FDR-corrected *P* value exceeding significance threshold of 0.05 are labelled with a star. The colours indicate super populations: oranges for African (**a** to **g**); greens for American (**h** to **k**); purples for East Asian (**l** to **p**); blues for European (**q** to **t**); pinks for South Asian (**u** to **y**).

**Extended Data Fig 2.**
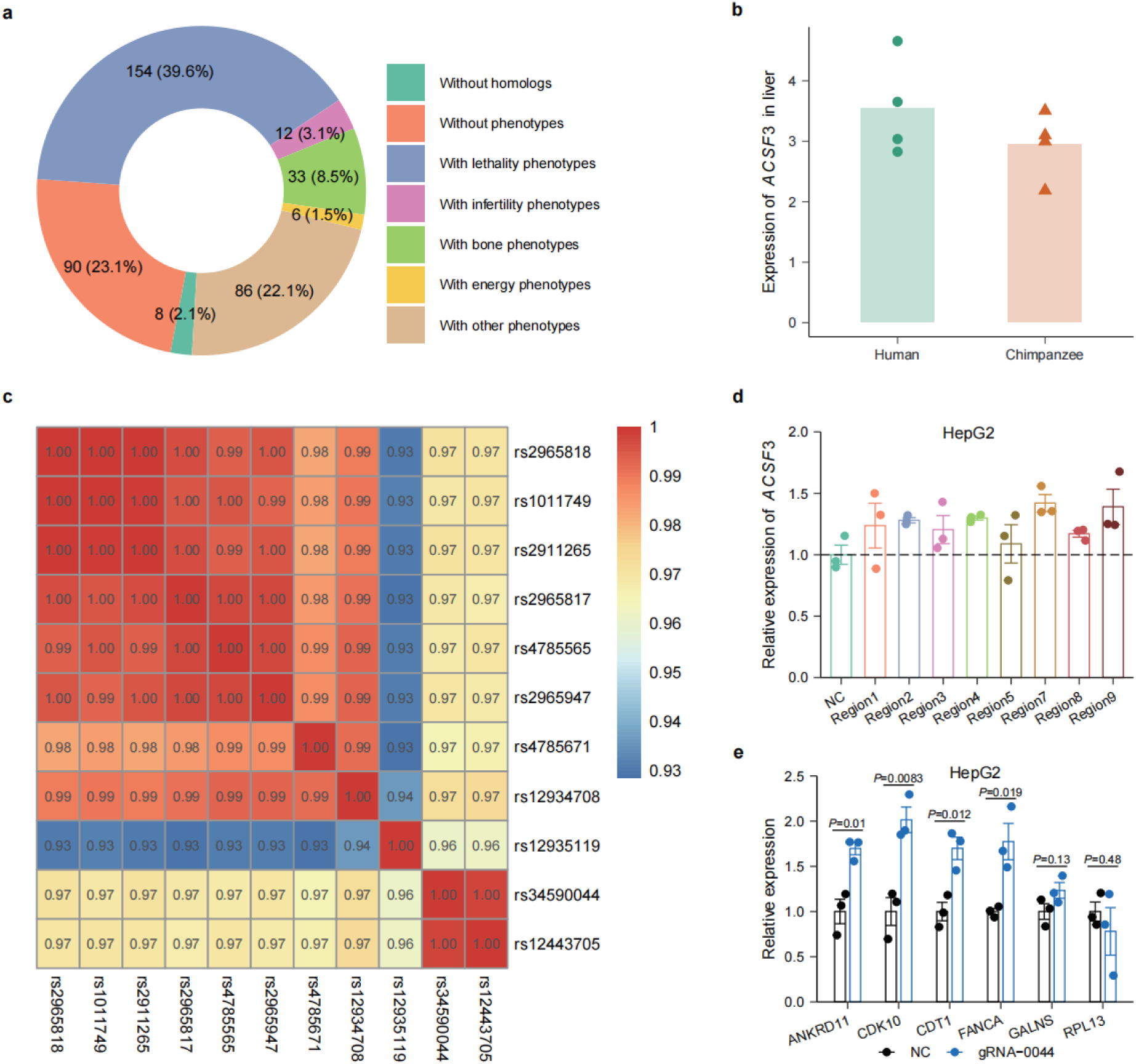
rs34590044-A is a representative AMH-specific mutation for height and BMR. **a**: The phenotypic categories of the 389 candidate genes likely to be affected by the height and BMR associated SNPs with AMH-specific alleles. The phenotypes of candidate genes in knocked-out mice are obtained from the Mouse Genome Informatics database. **b**: The expression of *ACSF3* in the liver of humans and chimpanzees. The y-axis shows both mean expression (bars) and expressions of each individual (dots; N=4 for each species). The data is obtained from Blake *et al*^1^. **c**: The linkage disequilibrium (R^2^) of 11 candidate causal SNPs of *ACSF3*. **d**: The expression of *ACSF3* upon the inhibition on the 10 SNPs (excluding rs34590044) with CRISPRi assay. N=3. **e**: The expression of the six neighbouring genes of *ACSF3* upon inhibiting rs34590044 with CRIPSRi assay. N=3.

**Extended Data Fig 3.**
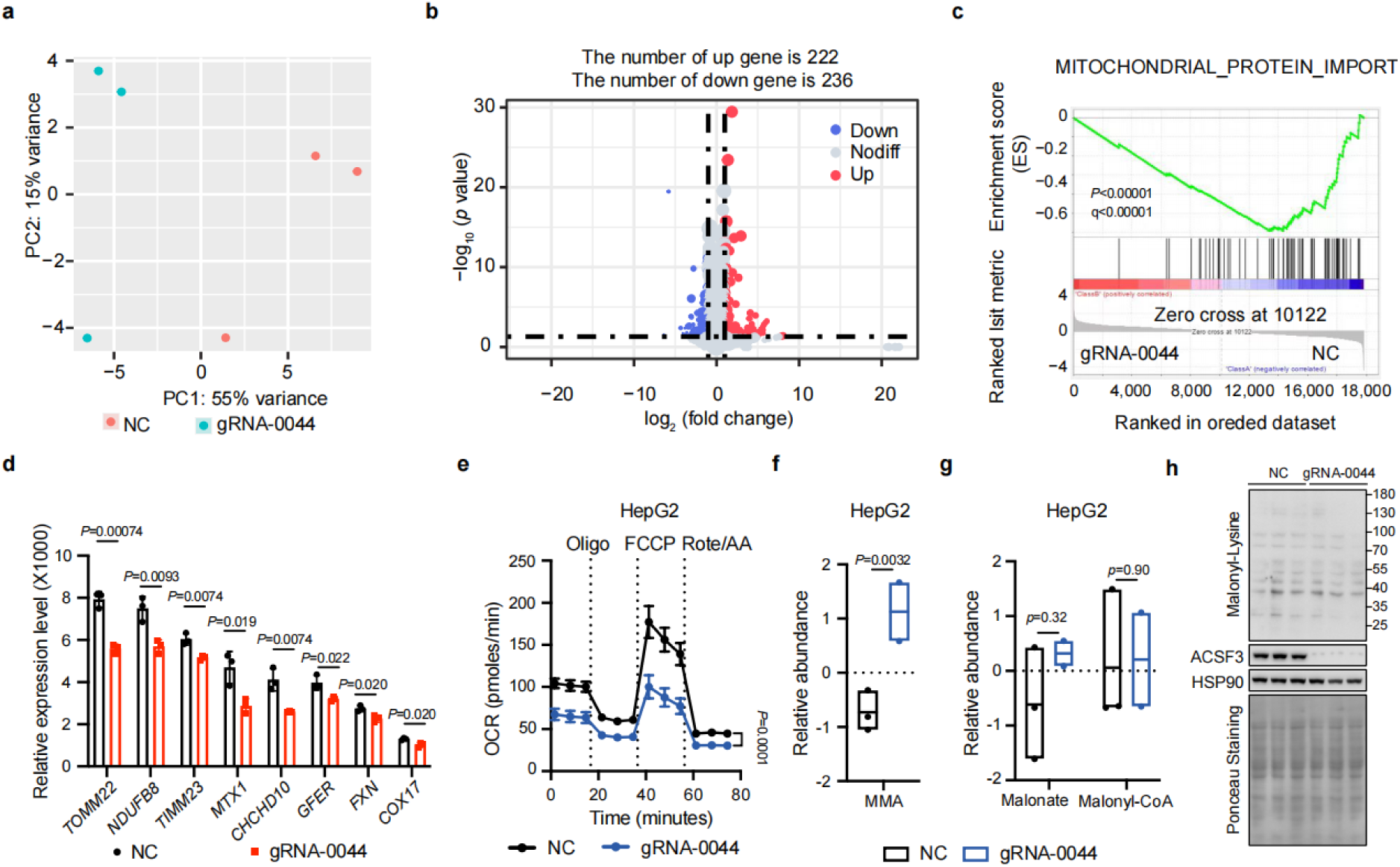
rs34590044-A regulates mitochondrial activity. **a-c**: RNA-seq analysis of HepG2 with inhibition on rs34590044 by CRISPRi. The results from Principal Component Analysis (**a**), Differentially Expression Analysis as volcano plot (**b**) and Gene Set Enrichment Analysis for mitochondrial activity related genes (**c**). NC: control. N=3. **d:** The expression of mitochondrial activity related genes in HepG2 cells with inhibition on rs34590044 by CRISPRi from RNA-seq analysis. NC: control. N=3. **e:** The respiratory rates of HepG2 cells with inhibition on rs34590044 by CRISPRi. N=3. **f-g:** Relative abundance of methylmalonate, malonate and malonyl-CoA in HepG2 cells with inhibition on rs34590044 by CRISPRi. NC: control. N_NC_=3. N_gRNA-0044_=2. **h:** The lysine malonylation of HepG2 with rs34590044-A CRISPRi. NC: control. N=3.

**Extended Data Fig 4.**
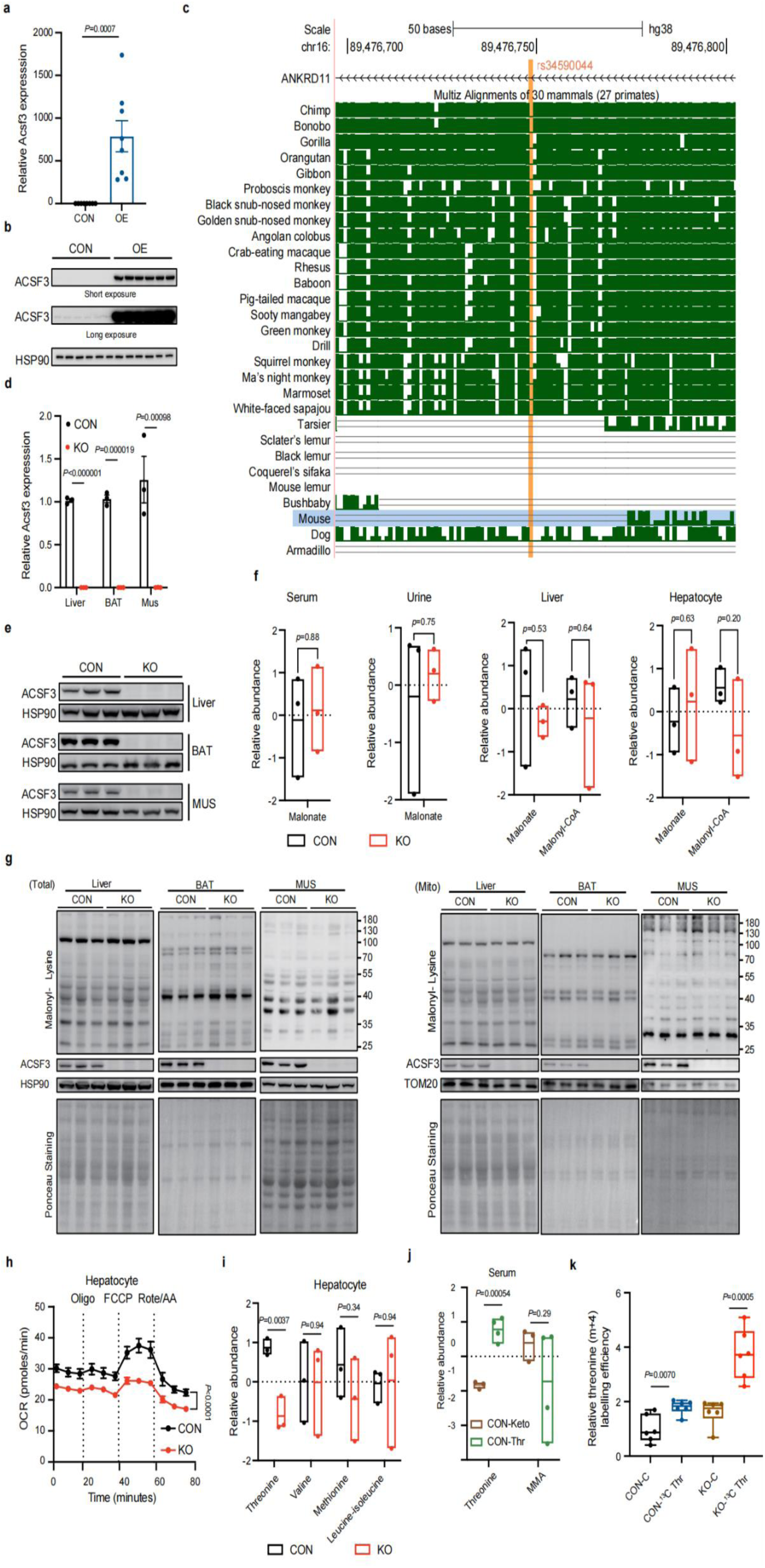
The overexpression and knock-out of *Acsf3* in mice. **a:** Relative expression of *Acsf3* mRNA in hepatocytes with *Acsf3* overexpression. CON: control. OE: overexpression. N=8. **b:** Relative protein expression of Acsf3 protein in hepatocytes with *Acsf3* overexpression via lenti-virus. CON: control. OE: overexpression. N=6. **c:** rs34590044 is located within a non-conserved region of the mouse genome. The figure is obtained from UCSC genome browser. **d-e:** Relative *Acsf3* mRNA (**d**) and Acsf3 protein (**e**) expression in liver, brown adipocytes (BAT) and muscle (MUS) with *Acsf3* knockout. CON: control mice. KO: knockout mice. N=3. **f:** Relative abundance of malonate and malonyl-CoA in serum, urine, liver and hepatocytes. CON: control mice. KO: knockout mice. N=3. **g:** The lysine malonylation of liver, brown adipocytes (BAT) and muscle (MUS) with *Acsf3* knockout. CON: control mice. KO: knockout mice. N=3. **h:** The respiratory rates in the hepatocytes of *Acsf3* KO mice. CON: control mice. KO: knockout mice. N=3. **i:** Relative abundance of threonine, valine, methionine and Leucine-isoleucine in hepatocytes of *Acsf3* KO mice. CON: control mice. KO: knockout mice. N=3. **j:** Relative abundance of threonine and methylmalonate in mice administered a ketogenic diet in drinking water. CON: control mice (*Acsf3^+/-^*). Keto: ketogenic diet. Thr: Threonine supplementary diet. MMA: methylmalonate. N=3-4. **k:** Relative abundance of ^13^C-labelled threonine in hepatocytes. CON: control mice (*Acsf3^+/-^*). KO: knockout mice (*Acsf3^-/-^*). C: chow diet. MMA-CoA: methylmalonyl-CoA. N=4.

**Extended Data Fig 5.**
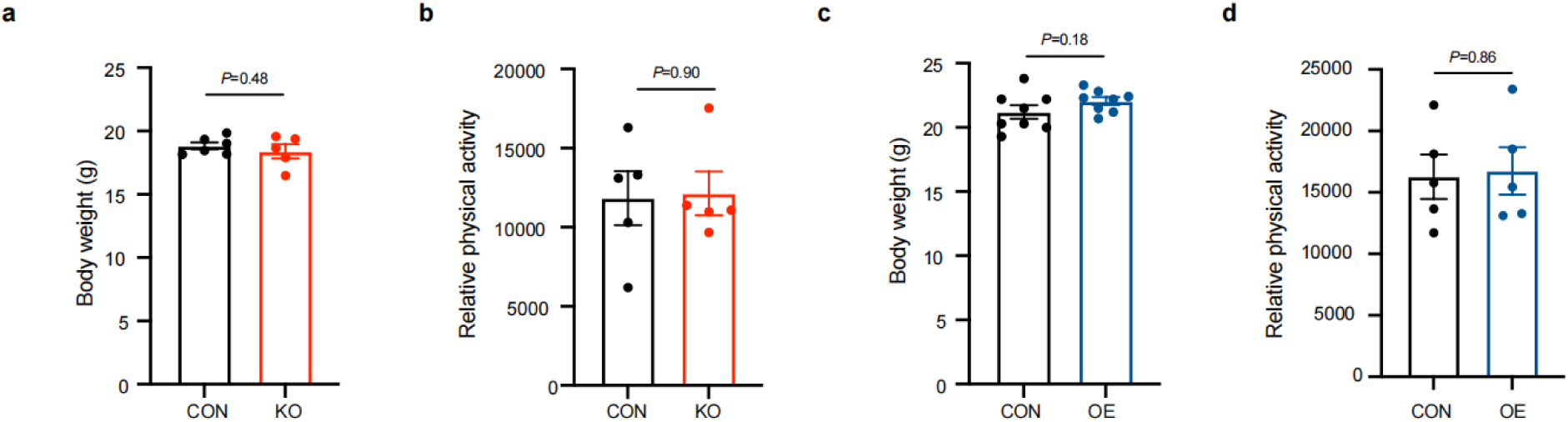
ACSF3 regulates body length and energy metabolism. **a:** The body weight in mice. CON: control mice (*Acsf3^+/-^*). KO: knockout mice (*Acsf3^-/-^*). N_CON_=6. N_KO_=5. **b:** Relative physical activity in mice. CON: control mice (*Acsf3^+/-^*). KO: knockout mice (*Acsf3^-/-^*). N=5. **c:** The body weight in mice. CON: control mice. OE: overexpression mice. N=8. **d:** Relative physical activity in mice. CON: control mice. OE: overexpression mice. N=5.

